# The scaffold protein PRR14L links the PP2A-TACC3 axis to mitotic fidelity and sensitivity to MPS1 inhibition

**DOI:** 10.1101/2025.11.04.686150

**Authors:** Albert Z. Liu, Akshay Narkar, Keming Li, Thierry Bertomeu, Blake A. Johnson, Jasmin Coulombe-Huntington, Yi Dong, Jin Zhu, Mike Tyers, Rong Li

## Abstract

Aneuploidy is a hallmark of cancer and is a potential vulnerability that can be selectively targeted. To systematically identify genes that affect the incidence and fitness of aneuploid cells, we conducted a genome-wide CRISPR/Cas9 screen using NMS-P715, an inhibitor of the spindle assembly checkpoint (SAC) kinase MPS1/TTK. In this study, we identified a number of genes known to regulate aneuploidy and mitosis, and subsequently focused on *PRR14L*, a ubiquitously expressed gene previously implicated in chronic myelomonocytic leukemia (CMML). Proximity labeling of PRR14L using TurboID revealed several cell division proteins, including the PP2A-B56 phosphatase complex and the spindle assembly factor TACC3, as PRR14L-interacting proteins. Loss of PRR14L prolongs SAC-dependent mitotic arrest in response to microtubule depolymerization but, paradoxically, leads to catastrophic mitotic errors upon SAC abrogation by MPS1 inhibitors. A model derived from our findings provides a rationale for exploiting MPS1 inhibition as a potential vulnerability in cancers containing either PRR14L loss of function mutations or FGFR-TACC3 fusions.

**Significance Statement:**

- Aneuploidy is a hallmark of cancer. Whether aneuploidy can be selectively targeted is not known.
- Utilizing a CRISPR/Cas9 screen, the authors found that loss of the gene PRR14L sensitizes cells to aneuploidy induction. Taking advantage of live cell imaging and proximity labeling, they linked PRR14L to TACC3 and mitosis.
- These findings suggest that spindle checkpoint inhibitors may have therapeutic potential in cancers with either loss-of-function PRR14L and/or gain of function TACC3 mutations.

## Introduction

Intratumor heterogeneity, drug resistance, and metastasis constitute major challenges in cancer therapy. All these have been associated with aneuploidy, a cellular state whereby the chromosome number is not a multiple of the haploid set. Aneuploidy in cancer is a consequence of chromosomal instability (CIN) due to heightened frequency of chromosome mis-segregation (Bakhoum and Landau, 2017; Bakhoum *et al*., 2018; Replogle *et al*., 2020). Common mechanisms that drive CIN in tumors include merotelic attachments, increased centrosome copy number, and whole genome duplication (Thompson and Compton, 2008; Gordon et al., 2012; Baker et al., 2024), but the molecular defects which underpin these mitotic errors remains incompletely understood. The spindle assembly checkpoint (SAC) is a key safeguard that prevents mitotic errors by delaying mitosis in the presence of improper kinetochore-microtubule attachments and other mitotic defects. Although mutations in SAC genes enhance tumorigenesis in animal models, SAC mutations are surprisingly rare in human tumors (Dobles *et al*., 2000; Baker *et al*., 2004; Schvartzman *et al*., 2010).

In this study, we performed a genome-wide CRISPR/Cas9 knockout (KO) screen in that aimed to identify genes and pathways that affect the frequency of or influence the cellular response to chromosomal mis-segregation. The screen was conducted with the MPS1/TTK inhibitor NMS-P715, and was designed to uncover genes whose loss either sensitized cells to SAC abrogation or modified the fitness of aneuploid progeny. Our screen identified *PRR14L,* a poorly understood nuclear gene which has loss of function mutations associated with clonality in chronic myelomonocytic leukemia (CMML) (Chase *et al*., 2019). In order to further characterize the function of *PRR14L*, we took advantage of proximity labeling with Turbo-ID to identify putative interacting proteins. The proximal proteome suggested that PRR14L may regulate mitosis, which we explored with live cell imaging in tandem with a variety of anti-mitotic treatments. Based on the putative link discovered between PRR14L and TACC3, we propose that MPS1/TTK inhibitors, such as NMS-P715, may have potentially unharnessed value as adjuvant treatments in tumors with either PRR14L loss of function and/or TACC3 gain of function mutations.

## Results and Discussion

### A CRISPR/Cas9 screen to identify enhancers and suppressors of MPS1i

To identify genes that may regulate aneuploidy induction and/or survival, we performed a genome-wide CRISPR/Cas9 KO screen utilizing a previously established NALM-6 pre-B cell lymphoblastic leukemia line, with a doxycycline-inducible Cas9, and the extended-knockout (EKO) sgRNA library harboring 278,754 sgRNAs (Bertomeu *et al*., 2018). Cells were treated for 7 days with doxycycline to induce Cas9 mediated gene KO, followed by a 2 day treatment with either 1 µM NMS-P715, to inhibit MPS1/TTK and induce aneuploidy, or vehicle control. NMS-P715 was subsequently washed out and cells were allowed to recover for 6 days in drug free media prior to analysis (Narkar *et al*., 2021). We then compared the frequency of sgRNAs between DMSO control and the NMS-P715 treated population to identify genes which may enhance or suppress the effect of MPS1/TTK inhibition (**Fig. 1A and B**).

**Figure 1.**
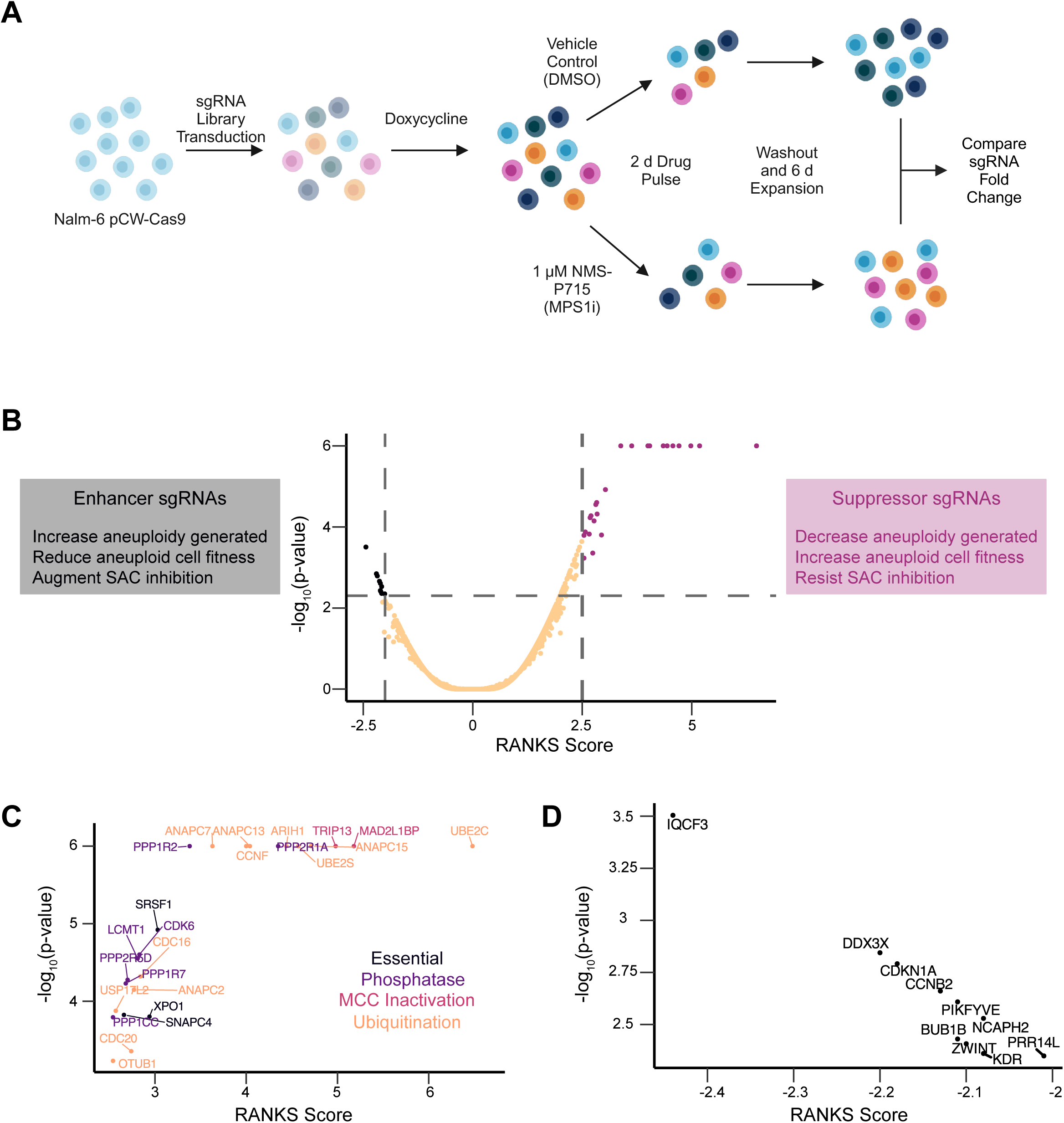
NALM-6 genome-wide CRISPR/Cas9 screen identifies *PRR14L* as an enhancer of NMS-P715. (A) Workflow of CRISPR/Cas9 screen (B) (left) Rationale underlying observation of enhancer sgRNAs (center) Volcano plot of CRISPR/Cas9 screen (right) Rationale underlying observation of suppressor sgRNAs. (C) Zoom-in on suppressor sgRNAs, with gene names highlighted, colored by gene category. (D) Zoom-in on enhancer sgRNAs, with gene names highlighted

Analysis of enriched sgRNAs revealed a number of genes that either encode anaphase-promoting complex (APC/C) subunits or regulate APC/C activity. This finding corroborates previous reports showing that a reduction in APC/C activity, which delays mitotic exit, can compensate for a disrupted SAC to limit chromosomal instability (**Fig. 1C**) (Sansregret *et al*., 2017). Another subset of suppressor genes encoded protein phosphatase 2A (PP2A) subunits or their regulators, consistent with the idea that loss of PP2A mitigates SAC silencing (Lara-Gonzalez *et al*., 2021).

The depleted sgRNAs identified by our screen affected a more diverse profile of genes (**Fig. 1d**). We identified a number of essential genes (*e.g. DDX3X and NCAPH2*), some of which also have well-established roles in mitosis (*e.g. BUB1B* and *ZWINT*). We decided to focus further investigation on *PRR14L*, a poorly understood gene hypothesized to play a role in cell division. *PRR14L* encodes a 237 kDa protein previously linked to myeloid neoplasms, PP2A, and PLK1 activity (Chase *et al*., 2019, 2023; Normandin *et al*., 2023).

### Proximity labeling to identify likely interacting proteins of PRR14L

To gain insight into the cellular function of *PRR14L*, we used TurboID (Branon *et al*., 2018; Cho *et al*., 2020) to identify the proximal proteome of PRR14L. HCT116 cells were generated which stably expressed PRR14L fused to 3xHA-TurboID via a glycine-serine linker (referred to as PRR14L-GS-3xHA-TurboID) and a spatial control for nuclear localization (3xHA-TurboID-3xNLS). We first validated nuclear localization of these construct with anti-HA antibodies and biotin labeling. In the spatial control cells with 3xHA-TurboID-3xNLS, we observed nuclear localization of the construct and strong labeling of the nucleoli. In contrast, PRR14L-GS-3xHA-TurboID localized primarily to the nucleoplasm and displayed weak biotin labeling of the nucleoplasm and nucleoli (**Fig. 2B**). To confirm that 3xHA-TurboID-3xNLS and PRR14L-GS-3xHA-TurboID labeled different proximal proteins, we generated cell lysates from untransfected, unlabeled and labeled cells and probed for both the HA-tag and biotinylated proteins (**Fig. 2C**). Western blot analysis revealed notably different biotin-labeling patterns between the two constructs, with strong PRR14L self-labeling and weaker labeling of other proteins.

**Figure 2.**
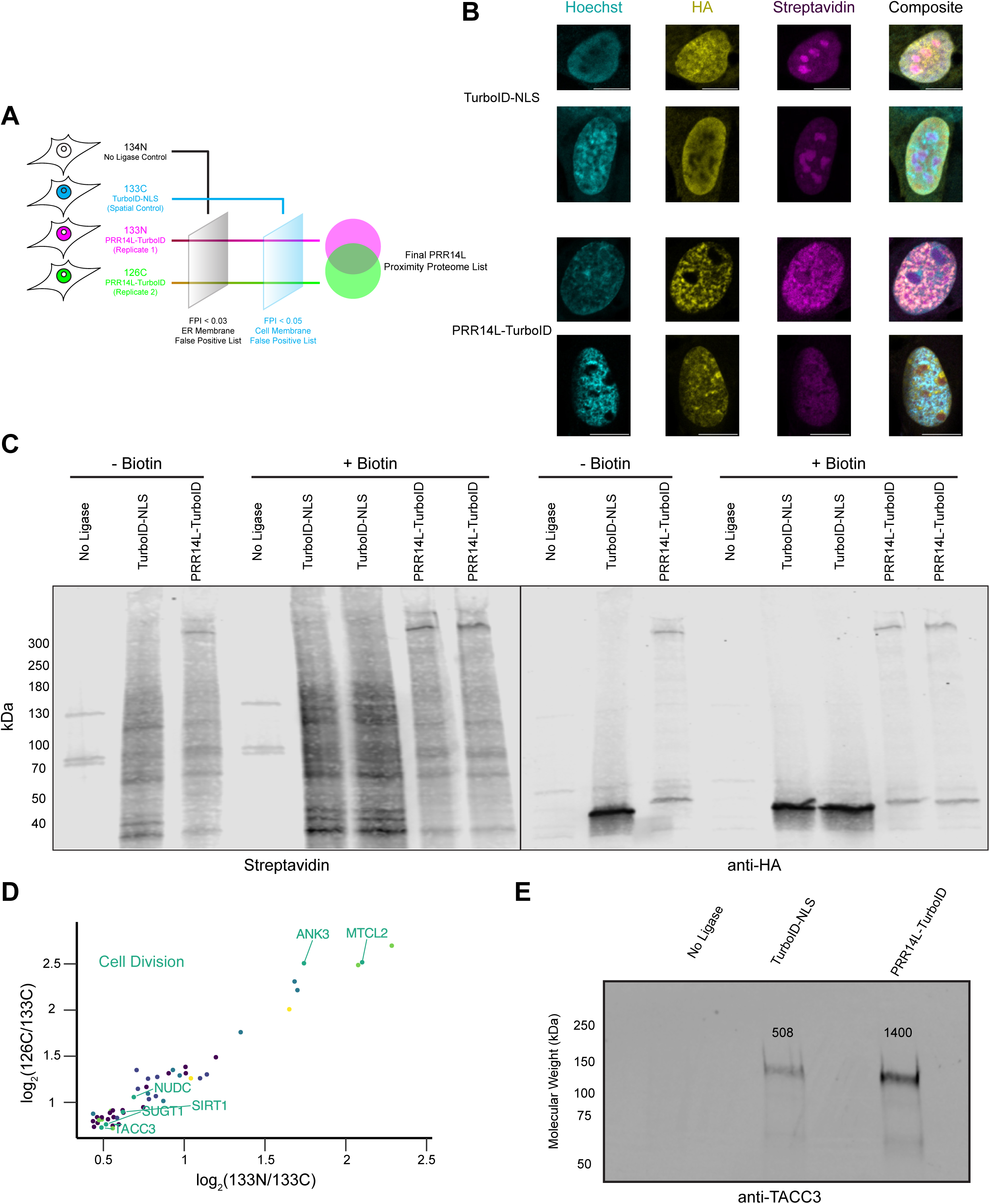
TurboID proximity labeling of PRR14L identifies multiple cell division proteins. (A) Schematic of TurboID Tandem Mass Tag (TMT) mass spectrometry experiment completed in HEK293T cells. (B) Differences between TurboID-NLS and PRR14L-TurboID localization. Representative images of hTERT RPE-1 cells transiently transfected with either 3xHA-TurboID-3xNLS or PRR14L-GS-3xHA-TurboID followed by 10 or 15 min labeling with 50 µM biotin respectively. Images acquired on LSM980, SR-8Y mode. (C) Characterization of biotinylation by PRR14L-TurboID. HEK293T cells were transiently transfected with either 3xHA-TurboID-3xNLS or PRR14L-GS-3xHA-TurboID followed with or without incubation with 50 µM biotin for 20 min. Cell lysates were used for western blot, and probed with anti-HA antibody and Streptavidin Dylight 680. Untransfected (no ligase) control also included for comparison. (D) PRR14L proximal proteome includes cell division proteins regulated by PP2A. Scatter plot of final PRR14L proximity proteome determined by mass spectrometry with PRR14L itself excluded. Dots colored by GO annotation. Proteins with GO annotations related to cell division have their names highlighted. (E) Western blot validation of PRR14L and TACC3 proximity. TurboID experiment was repeated, and cell lysates were utilized for anti-TACC3 western blot.

For mass spectrometry analysis, we proceeded with a 4-sample experiment design using a tandem mass tag (TMT) approach (**Fig. 2A**) (Branon *et al*., 2018). To control for endogenously biotinylated proteins and non-specific binding to the streptavidin beads, we included a no ligase control labeled with 134N. To account for background labeling of nuclear proteins, we included the spatial control, 3xHA-TurboID-NLS, labeled with 133C. Two independent experiments using PRR14L-GS-3xHA-TurboID, were labeled with 133N and 126C respectively (see Materials and Methods for further details).

We observed a linear correlation between labeling of the spatial control and both PRR14L independent experiments as expected (**Sup. Fig. 1A and B**). Importantly, the two independent experiments had a strong linear correlation with a R^2^ of 0.899 (**Sup. Fig. 1C**). We utilized ER membrane and cell membrane proteins as false positive filters (**Sup. Fig. 1D-G**, see methods). The two sets of filters reduced the potential PRR14L proximal proteome roughly 50-fold (**Sup. Fig. 1H and I**). Overlapping of the two independent experiments yielded the final PRR14L proximal proteome list, where PRR14L itself was the most enriched as expected (**Sup. Fig. 1J**). GO annotation identified several categories of proteins, including protein quality control, nuclear import, and mitochondria (**Sup. Fig. 1K-N**). A subset of proteins found were related to cell division and dephosphorylation (**Fig. 2D, Sup. Fig. 1K**). PRR14L has a C-terminal tantalus domain that is suggested to bind B56-family regulatory subunit of PP2A, and two of these subunits were identified within our Turbo-ID experiment (Serine/threonine-protein phosphatase 2A 56 kDa regulatory subunit alpha and delta isoforms). PRR14L proximity labeling also identified TACC3, which was validated via western blot (**Fig. 2E**). TACC3 is a centrosome and spindle protein that is a substrate of AURKA and thus hypothesized to be regulated by PP2A, as well as a known oncoprotein (Still *et al*., 1999; Lin *et al*., 2010; Wong *et al*., 2025).

#### Loss of PRR14L alters the response of cells to anti-mitotic drugs

To study the cellular function of PRR14L, we generated *PRR14L* knockouts (*PRR14L^-/-^*) in the near diploid HCT116 human colon carcinoma cell line. We chose this cell line because it is chromosomally stable and also adherent, allowing us to more easily visualize mitosis. The success of CRISPR knockout was validated via both genotyping and qPCR quantification of mRNA expression (**Sup. Fig. 2A and B**). Protein-level validation was not successful using commercially available antibodies for PRR14L, due to the presence of off-target antigens and likely also low endogenous expression of the protein (data not shown). As a non-targeting control (*NTC*), we generated monoclonal HCT116 cells transduced with LentiCRISPR v2 carrying a protospacer not predicted to be in the human genome.

Given the nuclear localization of PRR14L (Chase *et al*., 2019), we investigated whether *PRR14L^-/-^* caused a global change in gene transcription by bulk RNA-sequencing comparing *PRR14L^-/-^*cells with *NTC* cells. This revealed 375 differentially expressed genes and GO analysis suggested an effect on cell differentiation (Supplementary Tables 1,2). GSEA analysis revealed activation of interferon alpha response and epithelial mesenchymal transition signatures, as well as suppression of the G2/M checkpoint when comparing loss of PRR14L to control (**Sup Fig. 2C**). Further analysis with PROGENy revealed significant upregulation of the WNT pathway in PRR14L knockouts (**Sup Fig. 2D)**. However, the data does not suggest a general role for PRR14L in transcriptional regulation, and the differential gene expression that we observed could be attributable either to loss of PRR14L or cellular adaptation.

The proximity between PRR14L and both PP2A-B56 regulatory subunits and a putative PP2A substrate led us to hypothesize that PRR14L plays a role in regulating mitosis. To test this, we generated HCT116 *NTC* and HCT116 *PRR14L^-/-^* cells expressing both H2B-mNeonGreen to label chromatin, and mCherry-CAAX to label the cell membrane. Surprisingly, live cell imaging revealed no significant difference in NEBD to anaphase time between control and *PRR14L^-/-^*cells (**Fig. 3A**). Additionally, metaphase chromosome spreads revealed no significant difference in chromosome numbers between the control and *PRR14L^-/-^* cells (**Supplementary Fig. 3A**). These results suggested that under unperturbed conditions, *PRR14L^-/-^* cells undergo what appear to be normal mitoses.

**Figure 3.**
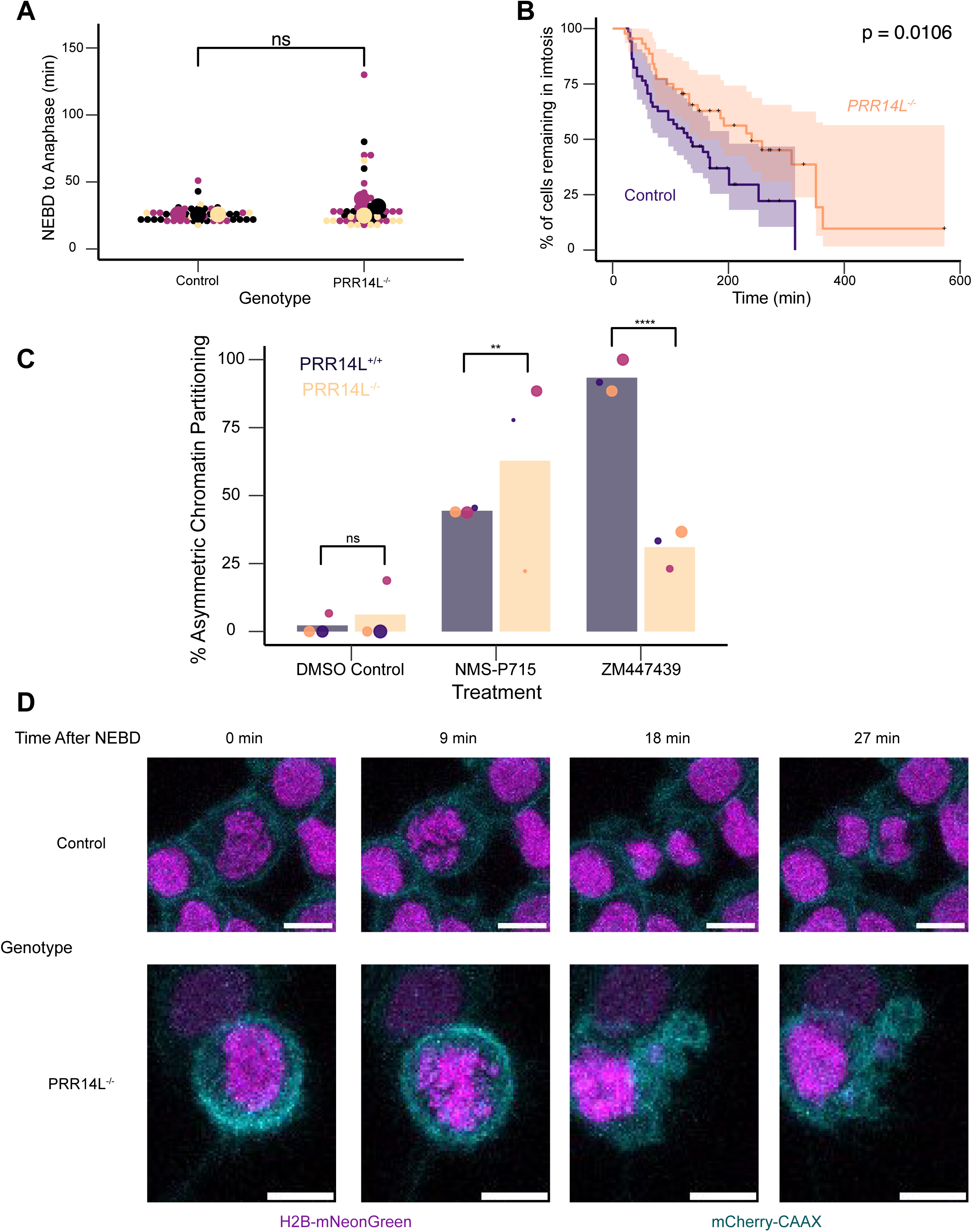
Loss of *PRR14L* alters the response to anti-mitotic drugs. (A) Quantification of untreated mitotic duration. Beeswarm plot comparing NEBD to anaphase time in HCT116 NTC and *PRR14L^-/-^* Clone A3 stably expressing H2B-mNeonGreen and mCherry-CAAX measured via live cell imaging. Colors denote independent experiments (n=3), small dots denote individual cells (NTC n = 48, *PRR14L^-/-^*n = 48), large dots denote mean of independent experiment. p-value calculated via Mann U-Whitney test using replicate averages, ns = non-significant. Images acquired on Zeiss LSM 780. (B) Quantification of mitotic duration in the presence of 25 nM nocodazole. Kaplan-Meier curve of % of cells still remaining in mitosis in HCT116 *NTC* and *PRR14L^-/-^*Clone A3 stably expressing H2B-mNeonGreen and mCherry-CAAX measured via live cell imaging. + marks denote censored events, where the cell still remained in mitosis at the end of the movie. n = 3 independent experiments. Total number of observed mitoses (NTC n = 51, PRR14L^-/-^ n = 44). p-value calculated via logrank test on combined data from independent experiments. Images acquired on Zeiss LSM 780. (C) Quantification of asymmetric chromosome partitioning in the presence of various drug treatments. Combined beeswarm and bar plot comparing proportion of cells undergoing asymmetric divisions between HCT116 *PRR14L^+/+^* and *PRR14L^-/-^*Clone A3 stably expressing H2B-mNeonGreen and mCherry-CAAX measured via live cell imaging. Colors denote independent experiments (n=3), dots denote mean of independent experiments with size correlated with number of mitoses observed (For DMSO, *PRR14L^+/+^* n = 72, *PRR14L^-/-^* n = 77. For 1 µM NS-P715, *PRR14L^+/+^* n = 68, *PRR14L^-/-^* n = 44. For 250 nM ZM447439 *PRR14L^+/+^* n = 68, *PRR14L^-/-^* n = 55). p-value calculated using two-proportion Z-test using combined data from independent experiments, ** = p < 0.01, **** = p < 0.0001. Images acquired on Zeiss LSM 780. (D) Representative images comparing live cell imaging of HCT116 *NTC* and *PRR14L^-/-^* Clone A3 stably expressing H2B-mNeonGreen and mCherry-CAAX with 1 µM NMS-P715 treatment. Z-max projections displayed. Frames synchronized to onset of NEBD. Scale bar width = 10 µm. Images acquired on Zeiss LSM 780.

Since PP2A-B56 is known to play a role in SAC silencing (Vallardi *et al*., 2019), we next tested the strength of the SAC in *PRR14L^-/-^* cells by treating cells with low dose (25 nM) nocodazole and observing the duration of mitosis. Notably, *PRR14L^-/-^* cells spent a significantly longer time in mitosis than the control in the presence of low dose nocodazole (**Fig. 3B**). This suggests that either PRR14L contributes to SAC silencing, and/or loss of PRR14L leads to mitotic defects that sensitize cells to nocodazole treatment. If the loss of PRR14L strengthened the SAC, then we would expect that *PRR14L^-/-^* cells should resist MPS1/TTK inhibition which weakens the SAC. However, *PRR14L*^-/-^ cells were found to have more prominent mitotic defects than control cells in the presence of NMS-P715 that frequently resulted in highly asymmetric chromatin distribution, with the most extreme case similar to monopolar division resulting in a single daughter cell with DNA inheritance (**Fig. 3C and D**). In line with this observation, metaphase chromosome spreads revealed that the ploidy of *PRR14L^-/-^* cells became near-tetraploid after NMS-P715 treatment (**Sup. Fig. 3B**).

It is possible that *PRR14L*^-/-^ cells only show mitotic defects when the SAC is disrupted by MPS1i (Colombo et al., 2010). We thus observed the response of PRR14L^-/-^ cells to ZM447439, an aurora kinase (AURK) inhibitor that disables SAC through an alternative mechanism. Surprisingly, we found that although control cells often fail cell division even with low-dose ZM447439 treatment, this cell division defect is partially suppressed by *PRR14L^-/-^* (**Fig. 3C, Sup. Fig. 3D**). Thus, loss of PRR14L appears to sensitize cells specifically to MPS1 inhibition and in fact compensates for AURK inhibition.

### Mutations affecting the PP2A-PRR14L-TACC3 axis sensitize cells to MPS1i

Given that PRR14L loss of function mutations have been found in CMML and are potentially sufficient for clonality, we asked whether MPS1i can be used to eliminate PRR14L deficient cells using a competition assay. HCT116-NTC cells were stably transfected with mNeonGreen and HCT116-*PRR14L^-/-^* with mScarlet. These cells were initially mixed with equal proportion and their relative abundance were tracked over 9 days in either DMSO or 1 µM NMS-P715 (**Fig. 4A and B**). *PRR14L^-/-^* cells were significantly more sensitive to MPS1i relative to control (**Fig. 4C**), and this finding also holds for both reversine and AZ-3146, other well-established MPS1/TTKi (**Sup. Fig. 3D**) (Hewitt *et al*., 2010; Santaguida *et al*., 2010).

**Figure 4.**
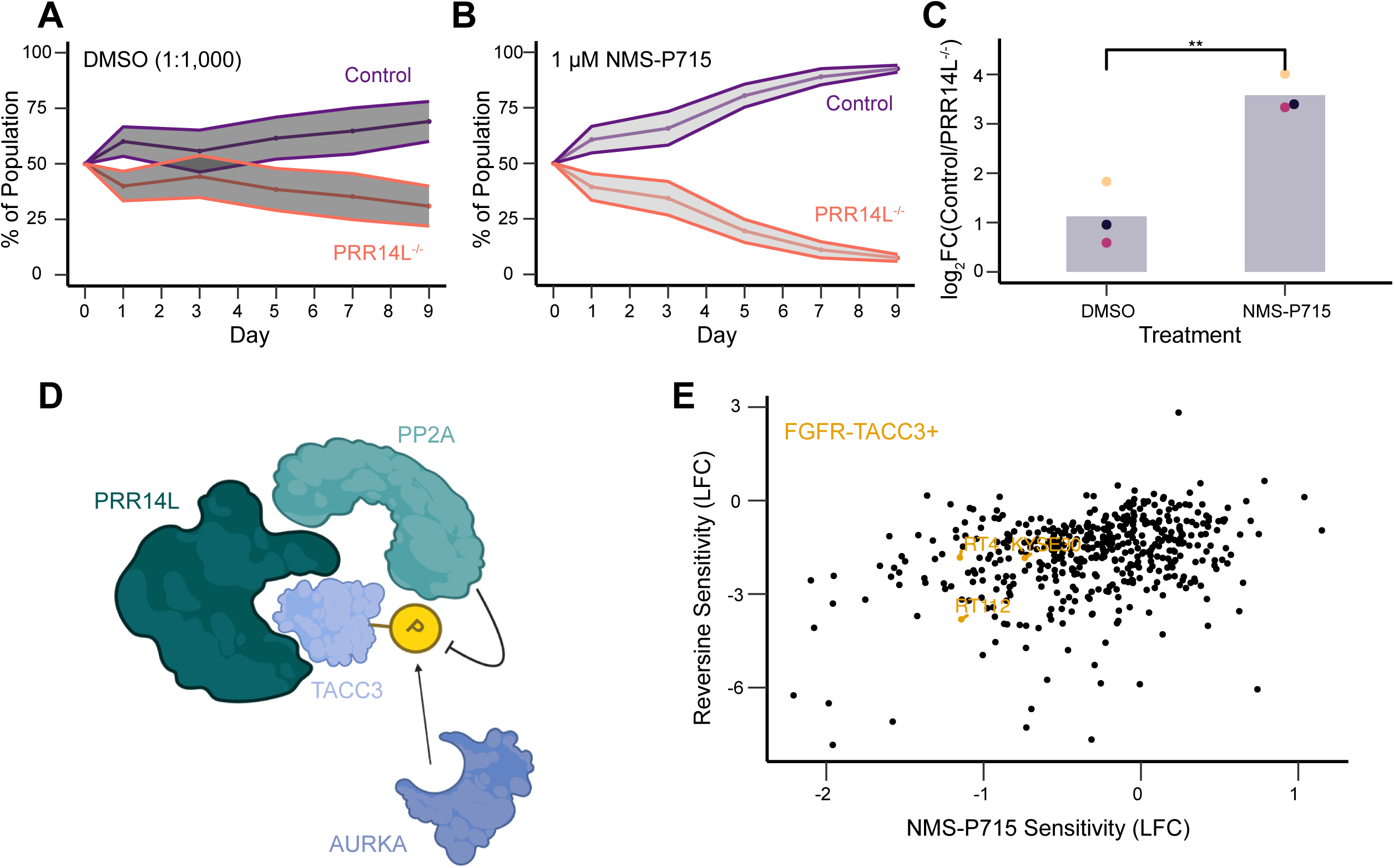
NMS-P715 has therapeutic potential for *PRR14L* loss of function and FGFR-TACC3 fusion tumors. (A) Flow cytometry tracking relative abundance of control and PRR14L^-/-^ cells over time. Cells were exposed to DMSO (1:1,000) during Days 1-3. Ribbons denote standard deviation, data shown from n = 2 independent experiments. (B) Flow cytometry tracking relative abundance of control and PRR14L^-/-^ cells over time. Cells were exposed to 1 µM NMS-P715 during Days 1-3. Ribbons denote standard deviation, data shown from n = 2 independent experiments. (C) Quantification of final timepoint (Day 9) of competition assay. Each dot represents a single independent experiment (n = 3), and each bar represents mean across independent experiments. P-value determined by one-sided student’s t-test, ** represents p < 0.01. (D) Working model for PRR14L regulation of TACC3, which may potentially explain *PRR14L^-/-^* phenotypes. (E) Comparison of tumor cell line sensitivity between NMS-P715 and Reversine. Each point represents a cancer cell line in the DepMap database. Three highlighted cell lines harbor FGFR-TACC3 fusions.

Based on findings described above, we propose a preliminary model whereby PRR14L serves as an adaptor protein that facilitates phosphatase access to a subset of its substrates (**Fig. 4D)**. This model posits that PRR14L acts as a scaffold bringing PP2A-B56 into proximity with TACC3, whose function is regulated by a phosphorylation balance between Aurora A kinase and PP2A. If true, this would suggest that loss of PRR14L could potentially phenocopy TACC3 overactivation due to FGFR-TACC3 gene fusions - driver mutations in cervical cancer, glioblastoma, and NSCLC (Singh *et al*., 2012; Capelletti *et al*., 2014; Carneiro *et al*., 2015). Both genetic alterations - loss of the PRR14L scaffold or FGFR-TACC3 fusion - may converge on TACC3 dysregulation, creating a synthetic lethal vulnerability to MPS1 inhibition. Supporting this idea, analysis of data from the Cancer Dependency Map portal (DepMap) revealed that the three cancer cell lines that harbor a FGFR-TACC3 fusion are sensitive to both NMS-P715 and reversine (**Fig. 4E**).

In summary, findings of this study provide further insight into the role of PRR14L as a regulator of mitotic fidelity whose function is particularly prominent when MPS1 is inhibited. We hypothesize that PRR14L functions as a scaffold for the PP2A-B56 phosphatase, coordinating the dephosphorylation of distinct substrates to ensure proper chromosome segregation. The paradoxical sensitivity to MPS1 and resistance to aurora kinase inhibition suggests a dual role for PRR14L. On one hand, its loss impairs PP2A-B56 mediated SAC silencing, leading to prolonged arrest in nocodazole. On the other hand, PRR14L appears critical for maintaining spindle integrity, likely via TACC3 regulation. When MPS1 is inhibited, the SAC is bypassed, and this underlying spindle fragility leads to catastrophic errors, overwhelming the cell despite the potentially hyperactive SAC machinery. Our results point to a potential vulnerability toward MPS1 inhibitors with driver mutations impact the PP2A-PRR14L-TACC3 axis and suggest a testable therapeutic strategy for such genetically defined cancers.

## Materials and Methods

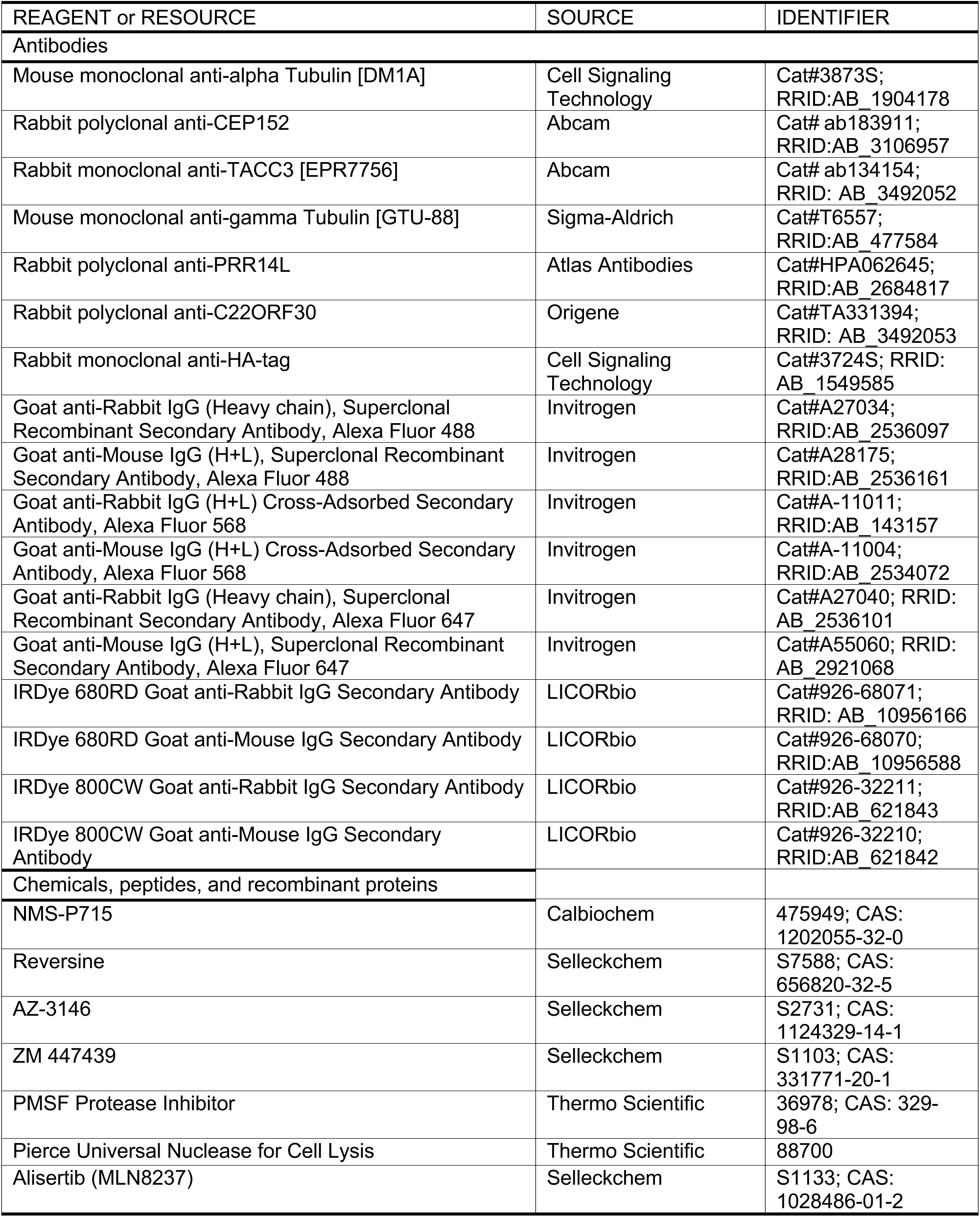
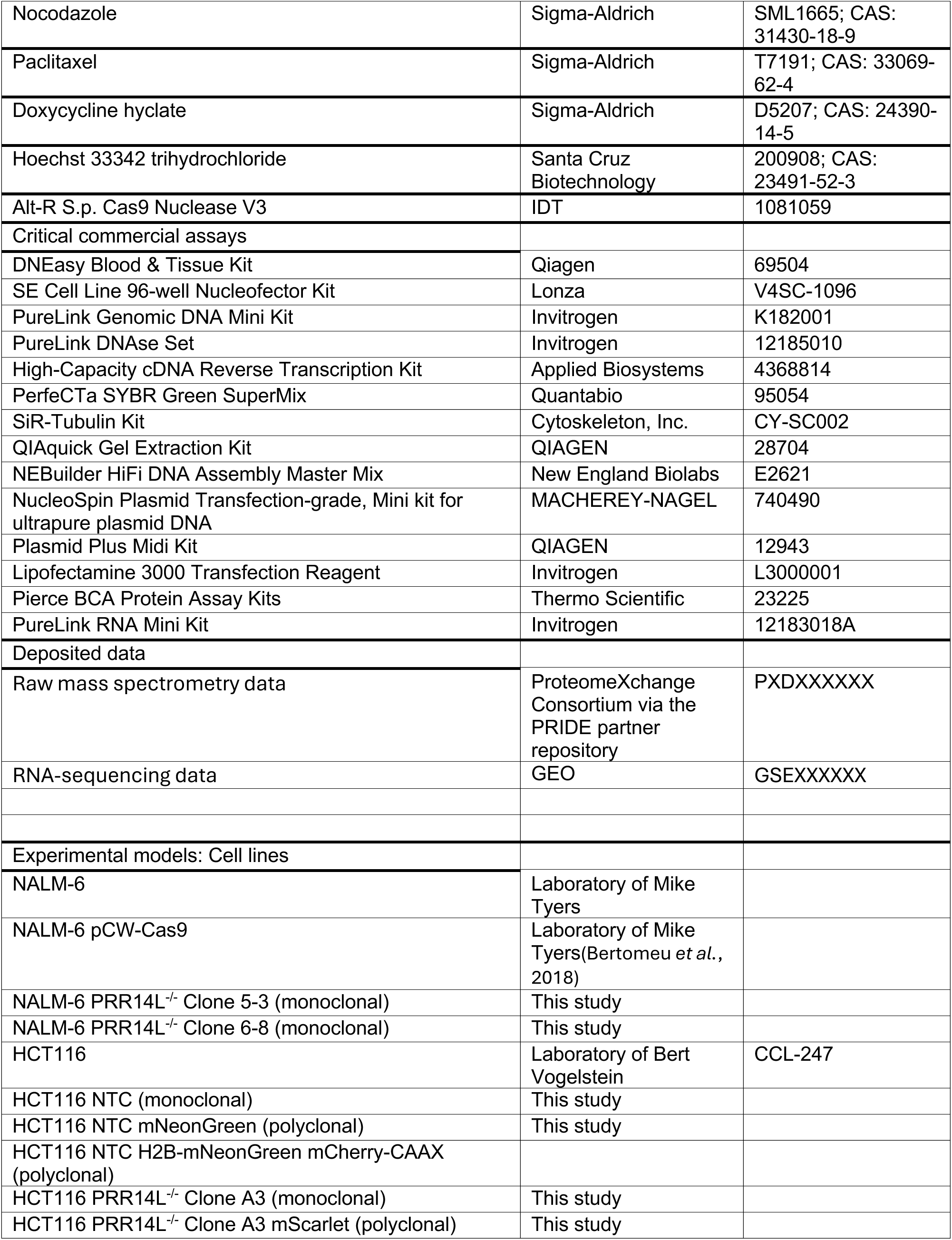
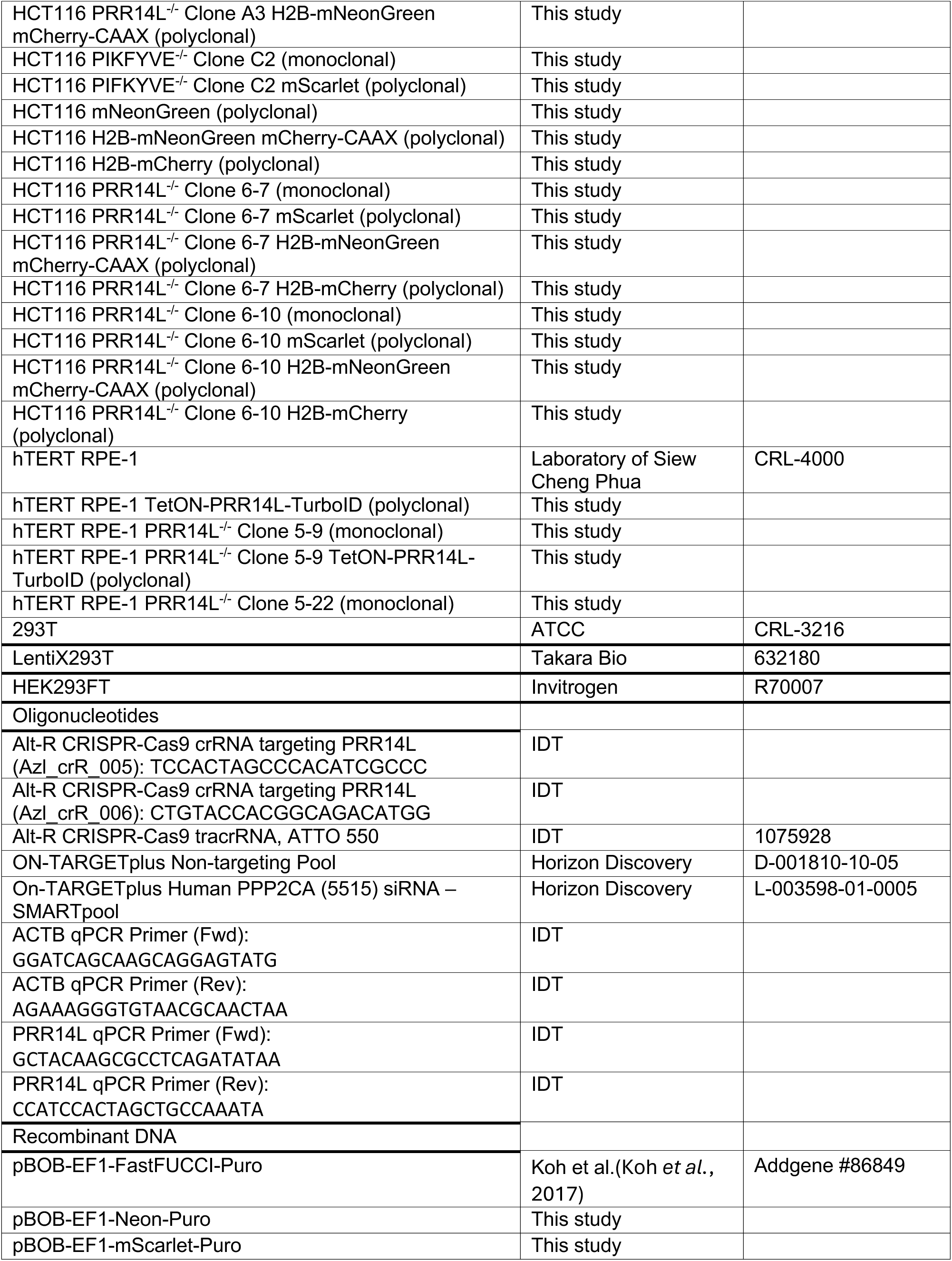
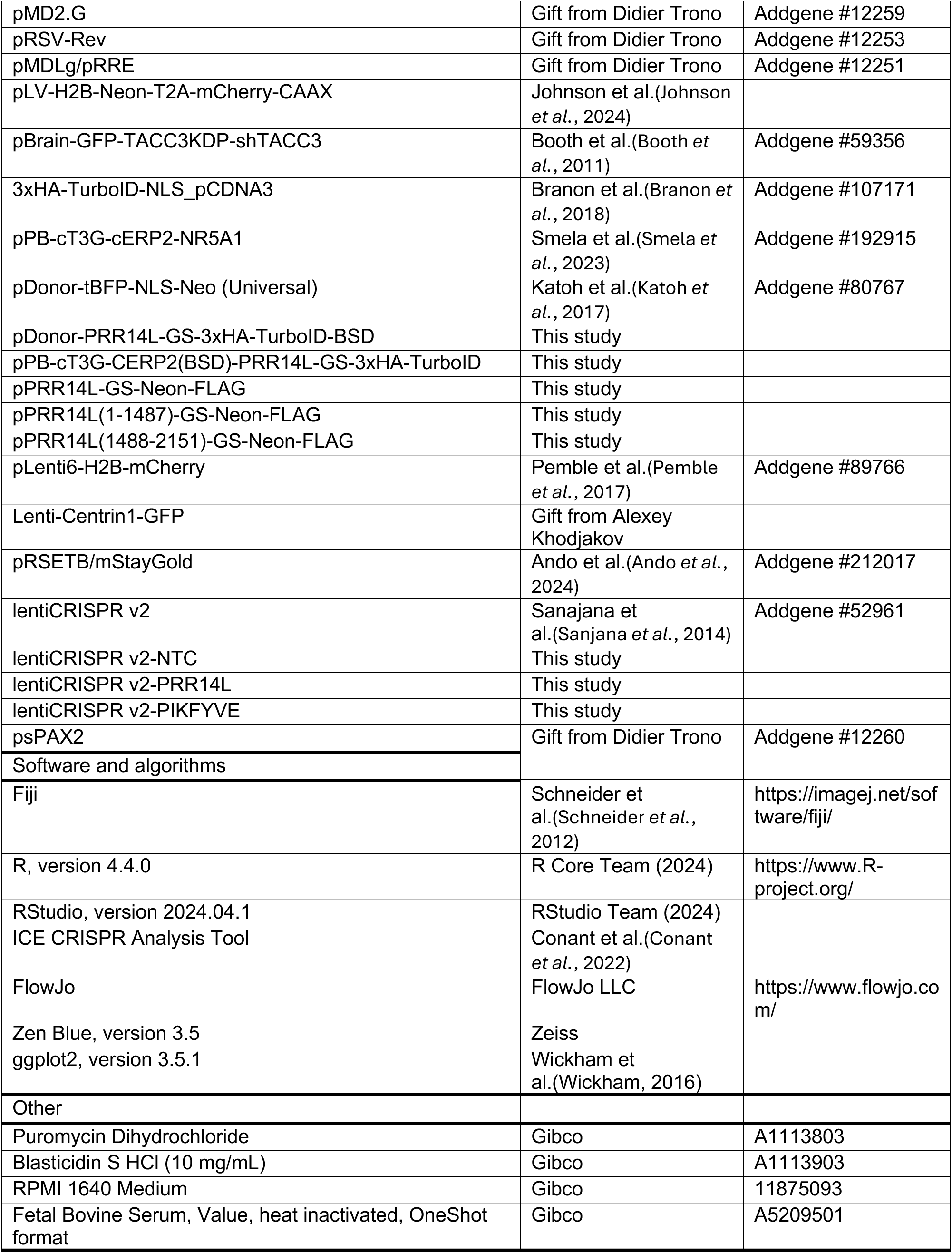
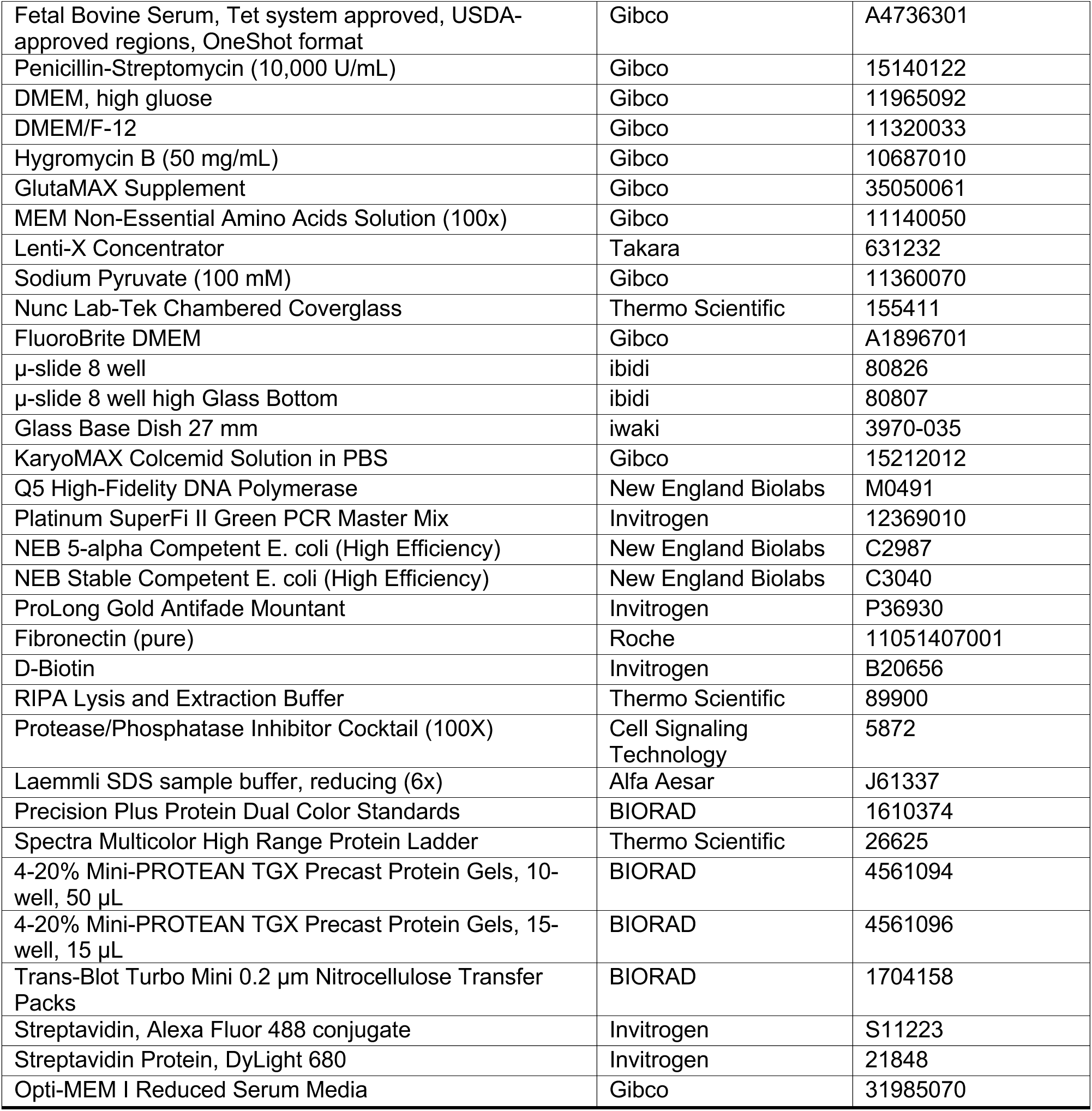

### Cell culture

NALM-6 cells were grown at 37 °C with 5% CO_2_ in RPMI 1640 supplemented with 10% heat-inactivated (HI) fetal bovine serum (FBS) and 100 U/mL Penicillin/Streptomycin (Pen/Strep). HCT116 cells were grown at 37 °C with 5% CO_2_ in DMEM supplemented with 10% HI FBS and 100 U/mL Pen/Strep. hTERT RPE-1 cells were grown at 37 °C with 5% CO_2_ in DMEM/F12 supplemented with 10% HI FBS and 10 µg/mL Hygromycin B. 293T cells were grown at 37 °C with 5% CO_2_ in DMEM supplemented with 10% HI FBS, 100 U/mL Pen/Strep, and 1% GlutaMAX. HEK293FT cells were grown at 37 °C with 5% CO_2_ in DMEM supplemented with 10% HI FBS. LentiX293T cells were grown at 37 °C with 5% CO_2_ in DMEM supplemented with 10% HI FBS, 1% GlutaMAX, 1x MEM non-essential amino acids, 1 mM Sodium Pyruvate, and 50 U/mL Pen/Strep. Mycoplasma testing was completed every 3-6 months to ensure cultures were free of mycoplasma contamination.

### Genome-wide pooled CRISPR/Cas9 Screen

The genome-wide pooled CRISPR/Cas9 screen was completed as previously described(Bertomeu *et al*., 2018; Normandin *et al*., 2023). In brief, a custom sgRNA library was used comprised of 233,268 sgRNAs that target 19,084 protein coding genes and 17,139 alternatively spliced exons. Each gene was targeted by approximately 10 sgRNAs. Noise was controlled for via 2,043 non-targeting sgRNAs. A doxycycline-inducible Cas9 clone of NALM-6 was used. The two conditions that were compared were vehicle control (DMSO, 1:1000) and 1 µM NMS-P715. After doxycycline induction for 1 week, DMSO and 1 µM NMS-P715 treatment were started Day 0 of the screen, and washed out Day 2. Cells were collected on both Day 2 and Day 8 for gDNA extraction and NGS.

### Generation of monoclonal KO cell lines

Two independent methods were used to generate monoclonal KO cell lines.

Lentivirus method:

HCT116 PRR14L^-/-^ clone A3 was generated using the lentiCRISPRv2 system. In brief, lentiCRISPRv2-PRR14L was cloned using the protospacer 5’-TCTGCTCTACCCTCTAATGG-3’, and lentiCRISPRv2-PIKFYVE was cloned using the protospacer 5’-ACATATGTTCGGTTAGCCTG-3’. Lentiviral particles were generated and collected via Lipofectamine-based transient transfection of HEK293FT cells with the following three plasmids in 1:1:1 molar ratio: lentiCRISPRv2, pMD2.G, and psPax2. HCT116 cells were transduced with lentiviral particles at a MOI of 0.5-1, and then underwent selection with 1 µg/mL Puromycin for 3 days followed by single cell sorting. Genomic DNA (gDNA) was extracted from single cell clones using DNEasy Blood and Tissue Kit (Qiagen) and amplified using the following primers: PRR14L: FWD 5’-CTGTTGCTACCATCCCCAGA-3’ | REV 5’-GAGACCACTTGAGGAAAGCCA-3’. PCR product was purified and submitted for sanger sequencing using the primer: 5’-CCACTTGAGGAAAGCCAGCT-3’. Sequencing data was analyzed using the ICE analysis tool to assess for successful gene knockout via frameshift mutation(Conant *et al*., 2022). Control cells referred to as “HCT116 NTC” were generated in the exact same manner except a non-targeting protospacer was used with the sequence 5’-gccccgccgccctcccctcc-3’.

### Lentivirus Generation and Transduction

Second generation lentivirus generation:

2 × 10^6^ HEK293FT cells were seeded in a T-25 flask. After 24 hrs, cells were transfected with the transfer plasmid, pMD2.G, and psPax2 in a 1:1:1 molar ratio using Lipofectamine 3000. 48 hrs after transfection, lentivirus supernatant was collected, aliquoted, and frozen at -80 °C for future use.

Third generation lentivirus generation:

Third generation lentivirus was produced following a previously published protocol with a few modifications25. LentiX293T cells were used for lentivirus production, and Lenti-X concentrator was used to concentrate lentiviral supernatants 5 to 10-fold. Concentrated lentiviral pellets were resuspended in DPBS and frozen at - 80 °C for future use.

Lentivirus transduction:

Lentivirus aliquots were warmed quickly using a 37 °C water bath for 2-3 min. Cells were transduced with a mixture of fresh media, lentivirus supernatant, and 10 µg/mL polybrene for 48 hrs. This was followed by either antibiotic selection or fluorescent-activated cell sorting (FACS) to obtain polyclonal and/or monoclonal cell lines.

### Competition Assay

Polyclonal fluorescent populations of competing cell lines were obtained by lentiviral transduction of cell lines with pBOB-EF1-Neon-Puro or pBOB-EF1-mScarlet-Puro. Cells were seeded at equal cell number on Day 0 in triplicate for each day to be measured, *i.e.*, for a 9-day experiment with two different drug treatments, a total of 30 wells were seeded initially. Media was changed to fresh media with drug treatment or control on Day 1. Drug treatments were washed out on Day 3. Cells were passaged every 2 days if the confluency was observed to be >75%; otherwise, a media change was completed.

Population numbers were measured using flow cytometry using either an Attune NxT Flow Cytometer or a Sony SH800. Data were analyzed using FlowJo and plots were generated with Rstudio and ggplot2.

### qPCR Profiling and Analysis

RNA was extracted from cells using PureLink RNA Mini Kit in combination with PureLink DNase Set. RNA concentration was measured with Nanodrop and then normalized to 100 ng/µL with RNase free water. cDNA was generated using High Capacity cDNA Reverse Transcriptase Kit. qPCR was completed using PerfeCTa SYBR Green Supermix. Fold change was calculated using ddCt method with ACTB and/or GAPDH as a control, and plots were generated with Rstudio and ggplot2.

### RNA Sequencing and Data Analysis

Library Preparation and Sequencing Total RNA was extracted from HCT116 NTC and PRR14L-/- (Clone A3) cells using the PureLink RNA Mini Kit (Invitrogen) with on-column DNase treatment (PureLink DNase Set, Invitrogen), according to the manufacturer’s instructions. Stranded RNA sequencing library preparation and high-throughput sequencing were performed by BGI Genomics (Hong Kong). Paired-end sequencing (PE100) was conducted on the DNBSEQ platform.

Preprocessing and Quantification Raw FASTQ files were processed by BGI using their internal analysis platform, Dr. Tom. Briefly, reads were quality-controlled, aligned to the human reference genome (Ensembl release 105), and gene expression levels were quantified as raw read counts.

Differential Gene Expression (DGE) Analysis Differential gene expression analysis was performed in R (v4.4.0) using the DESeq2 package (Love et al., 2014). To improve visualization and ranking, log2 fold changes (LFC) were shrunk using the apeglm method (Zhu et al., 2019). Genes were considered significantly differentially expressed if they met the criteria of an FDR-adjusted p-value (padj) < 0.05 and an absolute LFC > 1 (2-fold change). Gene Set Enrichment Analysis (GSEA) GSEA was performed using the clusterProfiler package (Wu et al., 2021) against the MSigDB Hallmark gene sets, retrieved via msigdbr. Genes were ranked by the Wald statistic obtained directly from the DESeq2 results. Pathway activities were inferred using PROGENy (Schubert et al., 2018). VST-normalized expression data, aggregated to gene symbols, were used as input and pathway scores were calculated based on the expression of the top 100 footprint genes per pathway. Differential pathway activities between KO and NTC samples were determined by applying a robust Student’s t-test to the inferred activity scores. Significance was determined by an FDR-adjusted p-value < 0.05.

### Live Cell Imaging

Live cell imaging was done via two methods based on location of experiment.

1. Cells were seeded in Nunc Lab-Tek chambered cover glass 2 days prior to imaging. Prior to the beginning of imaging, media was changed to fresh FluoroBrite DMEM supplemented with HI FBS and Pen/Strep, with or without drug treatment. Imaging was completed on a Zeiss LSM 780 laser scanning confocal microscope with an environmental chamber kept at 37 °C with 5% CO_2_ using a 63x oil-immersion objective (NA 1.4). For most independent experiments, 4-6 positions per genotype were imaged at once, with a 3 min time interval, and a 15-20 slice Z-stack with 1 µm slices. Deviations from this are mentioned in figure captions. Images were processed in ImageJ, with adjustments to color balance and cropping of selected cells of interest. Judgment of mitotic errors and/or mitotic outcome was done manually. Any fluorescence intensity quantification was done on unprocessed images.
2. Cells were seeded in ibidi µ-slides 2 days prior to imaging. 12 hrs prior to the start of imaging, media was changed to fresh FluoroBrite DMEM supplemented with HI FBS and 1x GlutaMAX. If drug treatment(s) were used, they were added 2 hrs prior to imaging in fresh FluoroBrite DMEM supplemented with HI FBS, 1x GlutaMax, and 50 nM SiR-Tubulin. Imaging was completed on one of three microscopes: Zeiss LSM 980 laser scanning confocal microscope with environmental chamber, 63x oil-immersion objective (NA 1.4), in either confocal mode, CO-8Y mode, or SR mode; Oxford Instruments BC43 spinning disk confocal microscope with environmental chamber, 60x oil-immersion objective (NA 1.42), and deconvolution completed with Imaris software; Zeiss Lattice Lightsheet 7 with environmental chamber, water-based objective (NA 1.0), and deskewing, cover glass transformation, and deconvolution was completed using Zen Blue software.

### Metaphase Spreads

Cells were treated with 100 ng/mL of colcemid for 6 hrs and then collected. Cell pellets were resuspended in PBS, and then incubated with pre-warmed 75 mM KCl for 10 min to induce cell swelling. Swollen cells were fixed with a solution of 3:1 methanol:glacial acetic acid overnight at 4 °C. Fixed cells were dropped onto pre-humidified glass slides, and then slides were dried by warming them to 65 °C on a hot plate for 20 min. Slides were mounted with ProLong Gold Antifade Mountant with DAPI followed by the addition of a coverslip and overnight curing at room temperature. Imaging was completed on a Nikon TiE-Eclipse epifluorescence microscope with 60x oil immersion objective. Chromosome number was recorded manually for each metaphase spread using the cell counter plugin in ImageJ.

### Plasmid cloning

A list of all plasmids used in this study can be found in the Recombinant DNA section of the materials list. All plasmid generation was completed via Gibson assembly (Gibson *et al*., 2009). Primers to generate new plasmids were designed using https://nebuilder.neb.com/<u>#!/</u>. For assemblies of 2 or 3 fragments, primers were designed with 20 bp overlaps. For assemblies of 4+ fragments, primers were designed with >24 bp overlaps. For assemblies requiring the addition of small fragments (e.g. GS linker, T2A sequence), small single-stranded oligonucleotide donors (ssODNs) were used. PCRs were completed using either Q5 Polymerase or Platinum SuperFi Polymerase. Products were run on 1% agarose gels, and then bands of the correct size were extracted using QIAquick Gel Extraction Kit. Gibson assembly was performed using NEBuilder HiFi DNA Assembly Master Mix, and products were transformed into either NEB 5-alpha Competent E. coli or NEB Stable Competent E. coli. Multiple single colonies were picked to inoculate miniprep cultures, and plasmid DNA was extracted using NucleoSpin Plasmid Transfection-grade, Mini kit. Plasmids were submitted for sanger sequencing to confirm assembly junctions. Confirmed plasmids were then re-picked to inoculate midiprep cultures, and plasmid DNA was extracted using QIAGEN Plasmid Plus Midi Kit. Midipreps were then submitted for sequencing of all functional genetic components (i.e. promoter, transgene, etc.) before being used for cell transfection.

### Transient Transfection

Transient transfection was carried out using Lipofectamine 3000, following the manufacturer’s instructions with a few modifications. The amount of OptiMEM Reduced Serum Media used was doubled, and prior to transfection the same amount of OptiMEM to be added was removed from cell culture. Lipofectamine 3000:P3000 Reagent::1.18:1. P3000 Reagent:Plasmid DNA::2 µL:1 µg. Cells were analyzed between 24-72 hrs after transfection.

### TurboID Immunofluorescence Validation

The following protocol is based on previously published work with slight modifications(Cho *et al*., 2020). Ibidi µ-slides 8-well were coated with 200 µL of 50 µg/mL fibronectin in DPBS per well for 30 min. 3.6×10^4^ 293T cells were seeded per well. After 24 hours, TurboID constructs were transfected using Lipofectamine 3000. After another 24 hours, media was replaced with fresh warm cDMEM supplemented with 50 µM biotin for either 10 min (3xHA-TurboID-3xNLS) or 20-30 min (PRR14L-GS-3xHA-TurboID). Labeling was stopped with 5x ice cold DPBS washes followed by fixation with 4% PFA in PBS at RT for 10-15 min. Blocking and permeabilization was completed simultaneously with 1 hr incubation with Abdil (2% BSA, 0.1% Triton X-100, in TBS).

1° antibody against HA-tag was incubated overnight (O/N) at 4 °C diluted 1:1,200 in Abdil. Cells were washed 3x with TBST (0.1% Triton X-100 in TBS) followed by 1hr room temperature (RT) incubation with 2° antibodies diluted in Abdil: anti-rabbit conjugated with Alexa Fluor 568 (1:1,000) and Streptavidin conjugated with Alexa Fluor 488 (1:1,000). Cells were washed 1x with TBST, followed by counterstaining with Hoechst 33342 in TBST for 5 min RT, followed by 1x TBST rinse, 1x TBST wash, and cells were imaged in 1x TBS buffer.

### Western Blot for TurboID Validation

3×10^5^ 293T cells were plated per well in 6-well plates. After 24 hours, TurboID constructs were transfected using Lipofectamine 3000. After another 24 hours, media was replaced with fresh warm cDMEM supplemented with 50 µM biotin for either 10 min (3xHA-TurboID-3xNLS) or 20-30 min (PRR14L-GS-3xHA-TurboID). Labeling was stopped with 5x ice cold DPBS washes followed by use of 1mL ice cold DPBS to detach cells and collect cell pellets. Cell pellets were resuspended in 100 µL of RIPA buffer supplemented with 1x protease and phosphatase inhibitor, 1 mM PMSF, and Pierce universal nuclease for cell lysis, and lysed for 20-40 min on ice with thorough pipetting every 10 min. Lysates were clarified by centrifugation at 14,000 rcf for 15 min at 4 °C. 80 µL of supernatant was collected and 5 µL was used for BCA protein quantification.

Protein concentration was normalized across samples and then combined with 6x Laemmli sample buffer and boiled for 10 min. Samples were loaded onto Mini-PROTEAN TGX precast protein gels, 4-20%, either 15×15 µL wells or 10×50 µL wells with either Precision Plus Dual Protein Color Standards or Spectra Multicolor High Range Protein Ladder for molecular weight ladder. Samples were transferred onto 0.2 µm nitrocellulose blots using the Trans-Blot Turbo transfer system, mixed molecular weight protocol. Blots were blocked with Intercept (TBS) Blocking buffer for 1 hr at RT followed by O/N incubation with 1° Ab (anti-HA). Blots were washed 4x with TBST (0.1% Tween-20 in TBS) followed by 1 hr incubation with 2° Ab diluted in 1:1::blocking buffer:TBST – Streptavidin Dylight 680 (1:5,000) and IRDye 800CW Goat anti-Rabbit IgG Secondary Antibody (1:20,000). Blots were washed again with 4x TBST followed by 2x TBS, and then imaged on an Odyssey CLx Imager.

### TMT Mass Spectrometry

2.25×10^6^ 293T cells were plated in T-75 flasks, two T-75 flasks per independent experiment. After 24 hours, TurboID constructs were transfected using Lipofectamine 3000. After another 24 hours, media was replaced with fresh warm cDMEM supplemented with 50 µM biotin for either 10 min (No ligase control and 3xHA-TurboID-3xNLS) or 30 min (PRR14L-GS-3xHA-TurboID). Labeling time of the PRR14L construct was extended to compensate for the weaker labeling by the PRR14L fusion construct relative to control. Labeling was stopped with 5x ice cold DPBS washes followed by use of 7.5 mL ice cold DPBS per flask to detach cells and collect cell pellets. Cell pellets were resuspended in 1.5 mL of RIPA buffer supplemented with 1x protease and phosphatase inhibitor, 1 mM PMSF, and Pierce universal nuclease for cell lysis, and lysed for 50 min on ice with thorough pipetting every 10 min. Lysates were clarified by centrifugation at 14,000 rcf for 15 min at 4 °C. Streptavidin magnetic beads were washed twice with 1 mL RIPA. 15 µL of lysate was retained as input, and the remainder was loaded on the streptavidin beads O/N rotating at 4 °C. 15 µL of flowthrough was collected per sample, and then beads were washed with: 2x 1 mL RIPA (2 min per wash), 1x 1mL KCl (2 min), 1x 1mL 100 nM Na_2_CO_3_ (10 sec), 1x 1mL 2M urea in 10 mM Tris-HCl pH 8.0 (10 sec), 2x 1 mL RIPA (2x 2min). The beads were resuspended in 1mL RIPA, transferred to fresh microcentrifuge tubes, and then flash frozen in liquid nitrogen and stored at -80 °C until further processing.

For each sample 100 µg of protein was added to a 10 kDa ultrafiltration tube, and volume was normalized across samples. Samples were centrifuged at 20 °C, 12,000 rcf for 20 min. 100 µL of 0.5M TEAB was added to samples and then centrifuged at 20 °C 12,000 rcf for 20 min (3x). Samples were subjected to Trypsin digestion at 37 °C for 4 hours. Peptide solution was concentrated and desalted, and the collected peptide fractions were freeze-dried. For peptide labeling, each tube of TMT (0.8 mg) was dissolved in 41 µL acetonitrile and shaken for more than 1 min. Peptide fragments were resuspended in 0.1M TEAB to a concentration of 3.74 µg/µL. 100 µg of peptide was labeled with 41 µL of TMT reagent for 2 hr at room temperature, ensuring pH was between 7-8.

For peptide fractionation the Shimadzu LC-20AB liquid phase system was used with a 5 µm 4.6×250mm Gemini C18 column for liquid phase separation of the sample. Dried peptide samples were reconstituted with mobile phase A (5% acetonitrile pH 9.8) and injected, eluting at a flow rate of 1 mL/min with the following gradients: 5% mobile phase B (95% acetonitrile, pH 9.8) for 10 min. 5% to 35% mobile phase B for 40 min. 35% to 95% mobile phase B for 1 min, mobile phase B for 3 min, and 5% mobile phase B for 10 min. The elution peak was monitored at a wavelength of 214 nm and one component was collected per minute, and the samples were combined according to the chromatography elution peak map to obtain 20 fractions, which were then freeze-dried.

For HPLC the dried peptide samples were reconstituted with mobile phase A (2% acetonitrile, 0.1% formic acid), centrifuged at 20,000 rcf for 10 min, and the supernatant was used for injection. Separation was performed using a Easy-nLC 1200, followed by a tandem self-packed C18 column (75 µm internal diameter, 1.9 µm column size, 25 cm column length) and separated at a flow rate of 200 nL/min using the following gradient: 0 – 3 min, 5% mobile phase B (80% acetonitrile, 0.1% formic acid); 3 – 45 min, mobile phase B was linearly increased from 8 to 44%; 45-50 min, mobile phase B increased from 44 to 60%; 50-53 min, mobile phase B rose from 60 to 100%; 53-60 min, 80% mobile phase B. The nanoliter liquid phase separation was directly connected to the mass spectrometer.

For mass spectrometry detection, the peptides separated by HPLC were ionized by a nanoESI source and then passed to a tandem mass spectrometer Orbitrap Exploris 480 for DDA mode detection. Parameters included: ion source voltage = 1.9 kV; MS1 mass spectrometer scanning range was 350-1,600m/z; resolution was set to 60,000; MS2 starting m/z was fixed at 100; resolution was 15,000. The ion screening conditions for MS2 fragmentation were: charge 2+ to 7+, and the first 30 parent ions with peak intensity exceeding 2×10^4^. Ion fragmentation mode was HCD, and the fragment ions were detected in Orbitrap. The dynamic exclusion time was set to 30 seconds. The automatic gain control was set to: MS1 300%, MS2 standard.

Mass spectrometry analysis was completed using Spectrum Mill MS Proteomics software as well as R.

To distinguish specific proximal proteins from abundant nuclear background, we employed a filtering strategy based on a custom False Positive Identification (FPI) rate, calculated against curated lists of proteins from irrelevant subcellular compartments (e.g., ER membrane, cell membrane) obtained from UniProt. This approach allowed us to apply stringent cutoffs to enrich for high-confidence interactors. A full list of identified proteins and quantitative values is provided in Supplementary Table 3. For the first filter where we compared abundance of proteins labeled by PRR14L relative to no ligase control, we applied a stringent cut-off of FPI < 0.03, given that we expect that almost all the proteins identified in our no ligase control should be non-specific, using a false positive list of ER membrane proteins. For the second filter where we compared abundance of proteins labeled PRR14L relative to the nuclear spatial control, a more relaxed cut-off of FPI < 0.05 was applied. This is because a greater percentage of labeling overlap is expected when both PRR14L and the spatial control occupy the nucleus. For this cut-off, we used a false positive list of cell membrane proteins.

## Acknowledgments

A.Z.L., and B.A.J. received support from the NIH Medical Scientist Training Program T32GM136577. This work was supported by Bloomberg Professorship funds to R.L. from Johns Hopkins University, a Mid-sized Grant (NRF-MSG-2023-0001) from Singapore National Research Foundation to R.L. and a Canadian Institutes of Health Research grant (FDN-167277) to M.T. We thank Angela Xiong for assistance with molecular cloning. Portions of the text were edited with the assistance of ChatGPT (OpenAI), a large language model AI tool.

## Data and code availability

Mass spectrometry and RNA-sequencing data will be deposited in publicly accessible databases prior to official publication.

## Figure Titles and Legends

**Figure S1.**

(A) Linear correlation between PRR14L-TurboID independent experiment 1 and TurboID-NLS. Scatter plot displaying all proteins with >2 unique peptides identified by mass spec, comparing replicate 1 against TurboID-NLS.

(B) Linear correlation between PRR14L-TurboID independent experiment 2 and TurboID-NLS. Scatter plot displaying all proteins with >2 unique peptides identified by mass spec, comparing replicate 2 against TurboID-NLS.

(C) Linear correlation between PRR14L-TurboID independent experiment 1 and independent experiment 2. Scatter plot displaying all proteins with >2 unique peptides identified by mass spec, comparing replicate 2 against replicate 1.

(D) Determination of filter 1 cut-off for independent experiment 1. False positive protein list used was ER membrane proteins from Uniprot. False positive identification plotted against log_2_ ratio comparing independent experiment 1 and no ligase control. Log_2_ cut-off determined at a FPI = 0.03 (3% of false positive proteins have log_2_ ratios higher than the cut-off of 0.592).

(E) Determination of filter 1 cut-off for independent experiment 2. False positive protein list used was ER membrane proteins from Uniprot. False positive identification plotted against log_2_ ratio comparing independent experiment 2 and no ligase control. Log_2_ cut-off determined at a FPI = 0.03 (3% of false positive proteins have log_2_ ratios higher than the cut-off of 0.804).

(F) Determination of filter 2 cut-off for independent experiment 1. False positive protein list used was cell membrane proteins from Uniprot. False positive identification plotted against log_2_ ratio comparing independent experiment 1 and spatial control. Log_2_ cut-off determined at a FPI = 0.05 (5% of false positive proteins have log_2_ ratios higher than the cut-off of 0.335).

(G) Determination of filter 2 cut-off for independent experiment 2. False positive protein list used was cell membrane proteins from Uniprot. False positive identification plotted against log_2_ ratio comparing independent experiment 2 and spatial control. Log_2_ cut-off determined at a FPI = 0.05 (5% of false positive proteins have log_2_ ratios higher than the cut-off of 0.603).

(H) Scatter plot displaying filter one and two for independent experiment 1.

(I) Scatter plot displaying filter one and two for independent experiment 2.

(J) Final PRR14L proximity proteome list. Most enriched protein is clearly PRR14L itself.

(K) Final PRR14L proximity proteome list with PRR14L excluded and phosphatase regulatory subunit genes highlighted, via GO annotations containing “dephosphorylation.”

(L) Final PRR14L proximity proteome list with PRR14L excluded and protein quality control genes highlighted, via GO annotations containing “proteasome,” “ubiquitin,” and/or “protein folding.”

(M) Final PRR14L proximity proteome list with PRR14L excluded and nuclear import genes highlighted, via GO annotation containing “import into nucleus.”

(N) Final PRR14L proximity proteome list with PRR14L excluded and mitochondrial genes highlighted, via GO annotation containing “mitoch.”

**Figure S2.**

(A) Genotyping validation of *PRR14L^-/-^* monoclonal HCT116 cells. Sanger sequencing traces aligned to genomic sequence from genome assembly hg38.

(B) qPCR quantification of PRR14L mRNA in *PRR14L^-/-^* HCT116 cells. mRNA fold change calculated via ddCt method, using *ACTB* as a control. Each dot represents a single independent experiment (n = 3), and each bar represents mean across independent experiments. P-value determined by one-sided student’s t-test, ** represents p < 0.01.

(C) Gene Set Enrichment Analysis (GSEA) of MSigDB Hallmark pathways. RNA-seq data from HCT116 *PRR14L^-/-^*(KO) and NTC cells were analyzed. Genes were ranked by the Wald statistic derived from DESeq2. The dot plot shows the significantly enriched pathways (FDR < 0.05), faceted by activation (positive Normalized Enrichment Score, NES) or suppression (negative NES) in *PRR14L^-/-^*cells. Dot size corresponds to the number of genes in the leading edge, and color indicates the FDR-adjusted p-value.

(A) (D) Differential pathway activity inference using PROGENy. VST-normalized expression data was used as input to estimate the activity of 14 major signaling pathways based on the expression of pathway footprint genes. The bar plot displays the change in activity score (KO - NTC). Bars are colored based on significance and direction of change. FDR values are indicated next to the corresponding bars.

**Figure S3.**

(A) Metaphase spread counts (untreated). Histogram comparing chromosome number between HCT116 NTC and HCT116 PRR14L^-/-^ Clone A3. Bin width displayed = 1. Data combined from n = 3 independent experiments. Total number of spreads counted (NTC n = 101, PRR14L^-/-^ n = 96). p-value calculated via two-sided student’s t-test using replicate averages.

(B) Metaphase spread counts (1 µM NMS-P715 treatment). Histogram comparing chromosome number between HCT116 NTC and HCT116 PRR14L^-/-^ Clone A3. Bin width displayed = 3. Data from n = 1 independent experiment. Total number of spreads counted (NTC n = 42, PRR14L^-/-^ n = 42).

(C) Representative images comparing live cell imaging of HCT116 NTC and PRR14L^-/-^ Clone A3 stably expressing H2B-mNeonGreen and mCherry-CAAX with 250 nM ZM 447439 treatment. Z-max projections displayed. Frames synchronized to onset of NEBD. Scale bar width = 10 µm. Images acquired on LSM780.

(D) Quantification of final timepoint (Day 9) of competition assay. Each dot represents a single independent experiment (n = 2 or 1), and each bar represents mean across independent experiments. DMSO values are the same as independent experiments 1 and 2 from Fig. 4C.

**Supplementary Table 3.** Full list of proteins identified and quantified in the PRR14L TurboID-MS experiment.

## References

Ando, R, Shimozono, S, Ago, H, Takagi, M, Sugiyama, M, Kurokawa, H, Hirano, M, Niino, Y, Ueno, G, Ishidate, F, et al. (2024). StayGold variants for molecular fusion and membrane-targeting applications. Nat Methods 21, 648–656.

Baker, DJ, Jeganathan, KB, Cameron, JD, Thompson, M, Juneja, S, Kopecka, A, Kumar, R, Jenkins, RB, De Groen, PC, Roche, P, et al. (2004). BubR1 insufficiency causes early onset of aging-associated phenotypes and infertility in mice. Nat Genet 36, 744–749.

Baker, TM, Waise, S, Tarabichi, M, and Van Loo, P (2024). Aneuploidy and complex genomic rearrangements in cancer evolution. Nat Cancer 5, 228–239.

Bakhoum, SF, and Landau, DA (2017). Chromosomal instability as a driver of tumor heterogeneity and evolution. Cold Spring Harb Perspect Med 7.

Bakhoum, SF, Ngo, B, Laughney, AM, Cavallo, JA, Murphy, CJ, Ly, P, Shah, P, Sriram, RK, Watkins, TBK, Taunk, NK, et al. (2018). Chromosomal instability drives metastasis through a cytosolic DNA response. Nature 553, 467–472.

Bertomeu, T, Coulombe-Huntington, J, Chatr-aryamontri, A, Bourdages, KG, Coyaud, E, Raught, B, Xia, Y, and Tyers, M (2018). A High-Resolution Genome-Wide CRISPR/Cas9 Viability Screen Reveals Structural Features and Contextual Diversity of the Human Cell-Essential Proteome. Mol Cell Biol 38, 1–24.

Booth, DG, Hood, FE, Prior, IA, and Royle, SJ (2011). A TACC3/ch-TOG/clathrin complex stabilises kinetochore fibres by inter-microtubule bridging. EMBO Journal 30, 906–919.

Branon, TC, Bosch, JA, Sanchez, AD, Udeshi, ND, Svinkina, T, Carr, SA, Feldman, JL, Perrimon, N, and Ting, AY (2018). Efficient proximity labeling in living cells and organisms with TurboID. Nat Biotechnol 36, 880–898.

Capelletti, M, Dodge, ME, Ercan, D, Hammerman, PS, Park, S Il, Kim, J, Sasaki, H, Jablons, DM, Lipson, D, Young, L, et al. (2014). Identification of recurrent FGFR3-TACC3 fusion oncogenes from lung adenocarcinoma. Clinical Cancer Research 20, 6551–6558.

Carneiro, BA, Elvin, JA, Kamath, SD, Ali, SM, Paintal, AS, Restrepo, A, Berry, E, Giles, FJ, and Johnson, ML (2015). FGFR3-TACC3: A novel gene fusion in cervical cancer. Gynecol Oncol Rep 13, 53–56.

Chase, A, Pellagatti, A, Singh, S, Score, J, Tapper, WJ, Lin, F, Hoade, Y, Bryant, C, Trim, N, Yip, BH, et al. (2019). PRR14L mutations are associated with chromosome 22 acquired uniparental disomy, age-related clonal hematopoiesis and myeloid neoplasia. Leukemia 33, 1184–1194.

Chase, A, Tarragona, GC, Lin, F, Yapp, S, Score, J, Bryant, C, and Cross, NCP (2023). Identification of B56 α , B56 γ and BAP1 as PRR14L binding partners. 1–15.

Cho, KF, Branon, TC, Udeshi, ND, Myers, SA, Carr, SA, and Ting, AY (2020). Proximity labeling in mammalian cells with TurboID and split-TurboID. Nat Protoc 15, 3971–3999.

Colombo, R, Caldarelli, M, Mennecozzi, M, Giorgini, ML, Sola, F, Cappella, P, Perrera, C, Re Depaolini, S, Rusconi, L, Cucchi, U, et al. (2010). Targeting the mitotic checkpoint for cancer therapy with NMS-P715, an inhibitor of MPS1 kinase. Cancer Res 70, 10255–10264.

Conant, D, Hsiau, T, Rossi, N, Oki, J, Maures, T, Waite, K, Yang, J, Joshi, S, Kelso, R, Holden, K, et al. (2022). Inference of CRISPR Edits from Sanger Trace Data. CRISPR Journal 5, 123–130.

Dobles, M, Liberal, V, Scott, ML, Benezra, R, and Sorger, PK (2000). Chromosome Missegregation and Apoptosis in Mice Lacking the Mitotic Checkpoint Protein Mad2, Li and Nicklas.

Gibson, DG, Young, L, Chuang, R-Y, Venter, JC, Hutchison, CA, and Smith, HO (2009). Enzymatic assembly of DNA molecules up to several hundred kilobases. Nat Methods 6, 343–345.

Gill, KP, and Denham, M (2020). Optimized Transgene Delivery Using Third-Generation Lentiviruses. Curr Protoc Mol Biol 133.

Gordon, DJ, Resio, B, and Pellman, D (2012). Causes and consequences of aneuploidy in cancer. Nat Rev Genet 13, 189–203.

Hewitt, L, Tighe, A, Santaguida, S, White, AM, Jones, CD, Musacchio, A, Green, S, and Taylor, SS (2010). Sustained Mps1 activity is required in mitosis to recruit O-Mad2 to the Mad1-C-Mad2 core complex. Journal of Cell Biology 190, 25–34.

Johnson, BA, Liu, AZ, Bi, T, Dong, Y, Li, T, Zhou, D, Narkar, A, Wu, Y, Sun, SX, Larman, TC, et al. (2024). Simple aneuploidy evades p53 surveillance and promotes niche factor-independent growth in human intestinal organoids. Mol Biol Cell.

Katoh, Y, Michisaka, S, Nozaki, S, Funabashi, T, Hirano, T, Takei, R, and Nakayama, K (2017). Practical method for targeted disruption of cilia-related genes by using CRISPR/Cas9-mediated, homology-independent knock-in system. Mol Biol Cell 28, 898–906.

Koh, S-B, Mascalchi, P, Rodriguez, E, Lin, Y, Jodrell, DI, Richards, FM, and Lyons, SK (2017). A quantitative FastFUCCI assay defines cell cycle dynamics at a single-cell level. J Cell Sci 130, 512–520.

Lara-Gonzalez, P, Pines, J, and Desai, A (2021). Spindle assembly checkpoint activation and silencing at kinetochores. Semin Cell Dev Biol 117, 86–98.

Lin, CH, Hu, CK, and Shih, HM (2010). Clathrin heavy chain mediates TACC3 targeting to mitotic spindles to ensure spindle stability. Journal of Cell Biology 189, 1097–1105.

Narkar, A, Johnson, BA, Bharne, P, Zhu, J, Padmanaban, V, Biswas, D, Fraser, A, Iglesias, PA, Ewald, AJ, and Li, R (2021). On the role of p53 in the cellular response to aneuploidy. Cell Rep 34, 108892.

Normandin, K, Coulombe-Huntington, J, St-Denis, C, Bernard, A, Bourouh, M, Bertomeu, T, Tyers, M, and Archambault, V (2023). Genetic enhancers of partial PLK1 inhibition reveal hypersensitivity to kinetochore perturbations. PLoS Genet 19.

Pemble, H, Kumar, P, van Haren, J, and Wittmann, T (2017). GSK3-mediated CLASP2 phosphorylation modulates kinetochore dynamics. J Cell Sci 130, 1404–1412.

Replogle, JM, Zhou, W, Amaro, AE, Mcfarland, JM, Villalobos-Ortiz, M, Ryan, J, Letai, A, Yilmaz, O, Sheltzer, J, Lippard, SJ, et al. (2020). Aneuploidy increases resistance to chemotherapeutics by antagonizing cell division.

Sanjana, NE, Shalem, O, and Zhang, F (2014). Improved vectors and genome-wide libraries for CRISPR screening. Nat Methods 11, 783–784.

Sansregret, L, Patterson, JO, Dewhurst, S, López-García, C, Koch, A, McGranahan, N, Chao, WCH, Barry, DJ, Rowan, A, Instrell, R, et al. (2017). APC/C dysfunction limits excessive cancer chromosomal instability. Cancer Discov 7, 218–233.

Santaguida, S, Tighe, A, D’Alise, AM, Taylor, SS, and Musacchio, A (2010). Dissecting the role of MPS1 in chromosome biorientation and the spindle checkpoint through the small molecule inhibitor reversine. Journal of Cell Biology 190, 73–87.

Schneider, CA, Rasband, WS, and Eliceiri, KW (2012). NIH Image to ImageJ: 25 years of image analysis. Nat Methods 9, 671–675.

Schvartzman, JM, Sotillo, R, and Benezra, R (2010). Mitotic chromosomal instability and cancer: Mouse modelling of the human disease. Nat Rev Cancer 10, 102–115.

Singh, D, Chan, JM, Zoppoli, P, Niola, F, Sullivan, R, Castano, A, Liu, EM, Reichel, J, Porrati, P, Pellegatta, S, et al. (2012). Transforming Fusions of FGFR and TACC Genes in Human Glioblastoma. Science (1979) 337, 1231–1235.

Smela, MDP, Kramme, CC, Fortuna, PRJ, Adams, JL, Su, R, Dong, E, Kobayashi, M, Brixi, G, Kavirayuni, VS, Tysinger, E, et al. (2023). Directed differentiation of human iPSCs to functional ovarian granulosa-like cells via transcription factor overexpression. Elife 12.

Still, IH, Vince, P, and Cowell, JK (1999). The Third Member of the Transforming Acidic Coiled Coil-Containing Gene Family, TACC3, Maps in 4p16, Close to Translocation Breakpoints in Multiple Myeloma, and Is Upregulated in Various Cancer Cell Lines.

Thompson, SL, and Compton, DA (2008). Examining the link between chromosomal instability and aneuploidy in human cells. Journal of Cell Biology 180, 665–672.

Vallardi, G, Allan, LA, Crozier, L, and Saurin, AT (2019). Division of labour between pp2a-b56 isoforms at the centromere and kinetochore. Elife 8, 1–25.

Wickham, H (2016). ggplot2 Elegant Graphics for Data Analysis Second Edition, Houston.

Wong, S-S, Monteiro, JM, Chang, C-C, Peng, M, Mohamed, N, Steinacker, TL, Xiao, B, Saurya, S, Wainman, A, and Raff, JW (2025). Centrioles generate two scaffolds with distinct biophysical properties to build mitotic centrosomes. Sci Adv.

