## Supplementary figures and images for "The scaffold protein PRR14L links the PP2A-TACC3 axis to mitotic fidelity and sensitivity to MPS1 inhibition"

### Supplemental Figures

#### Supplemental Figure 1

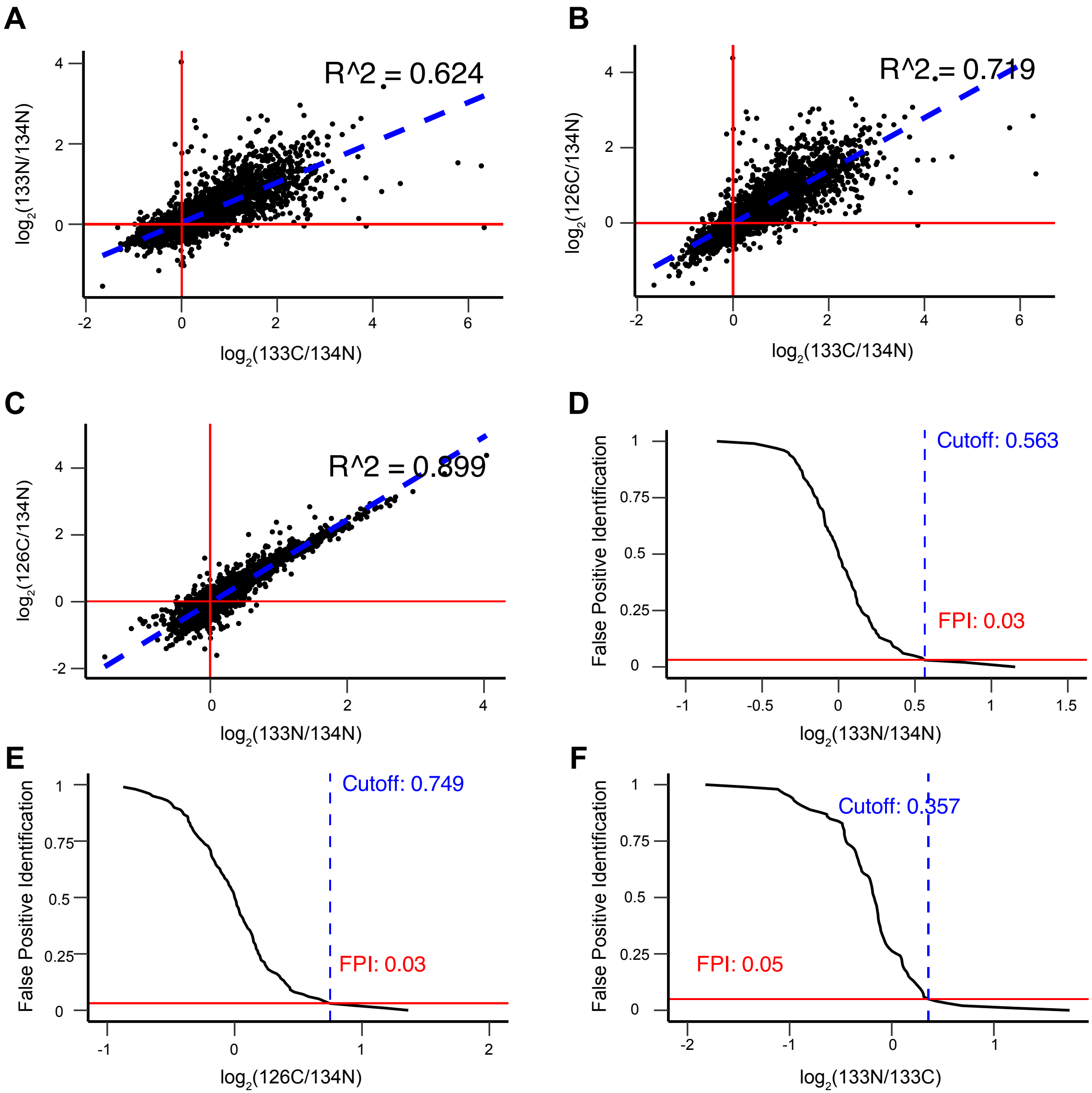

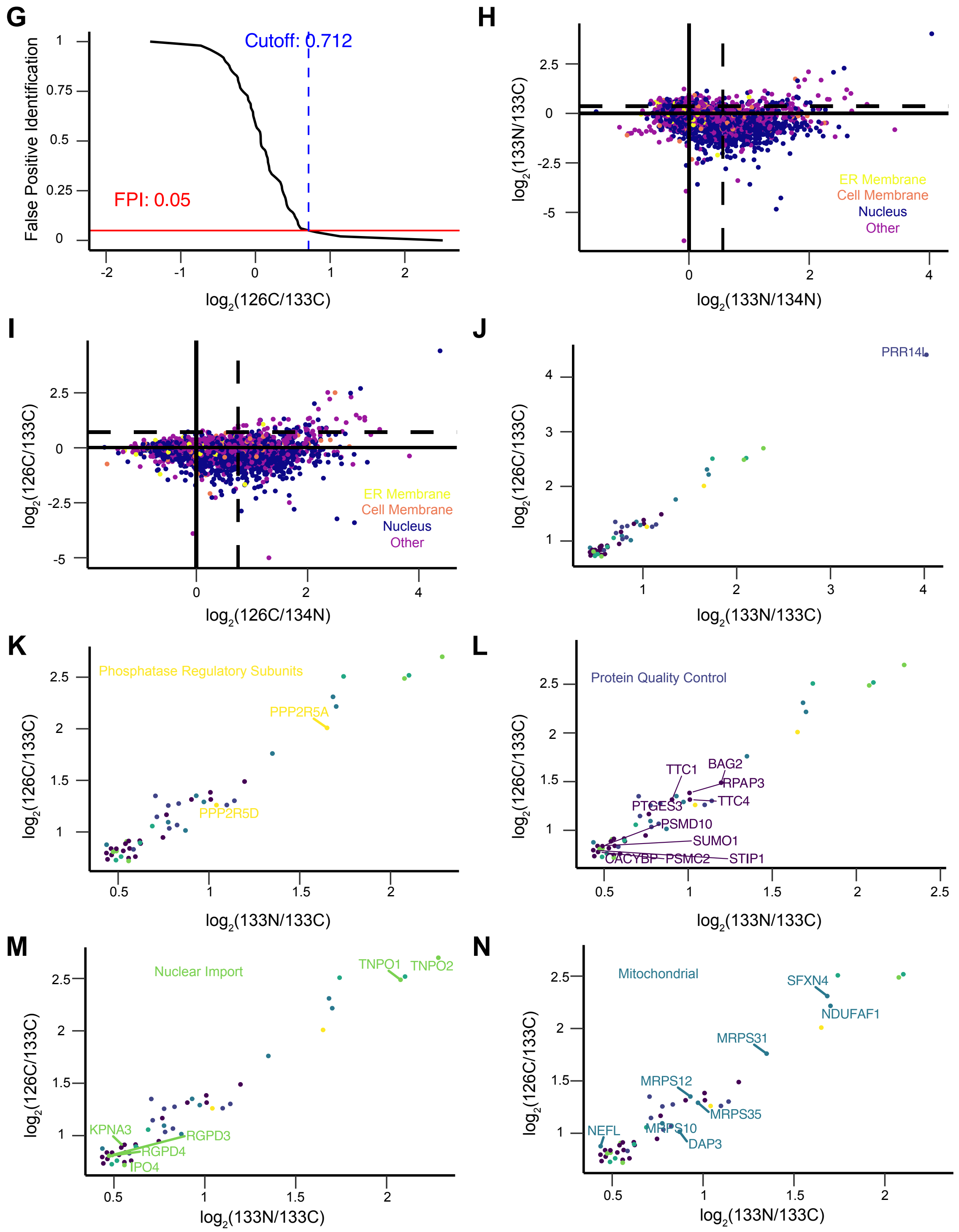

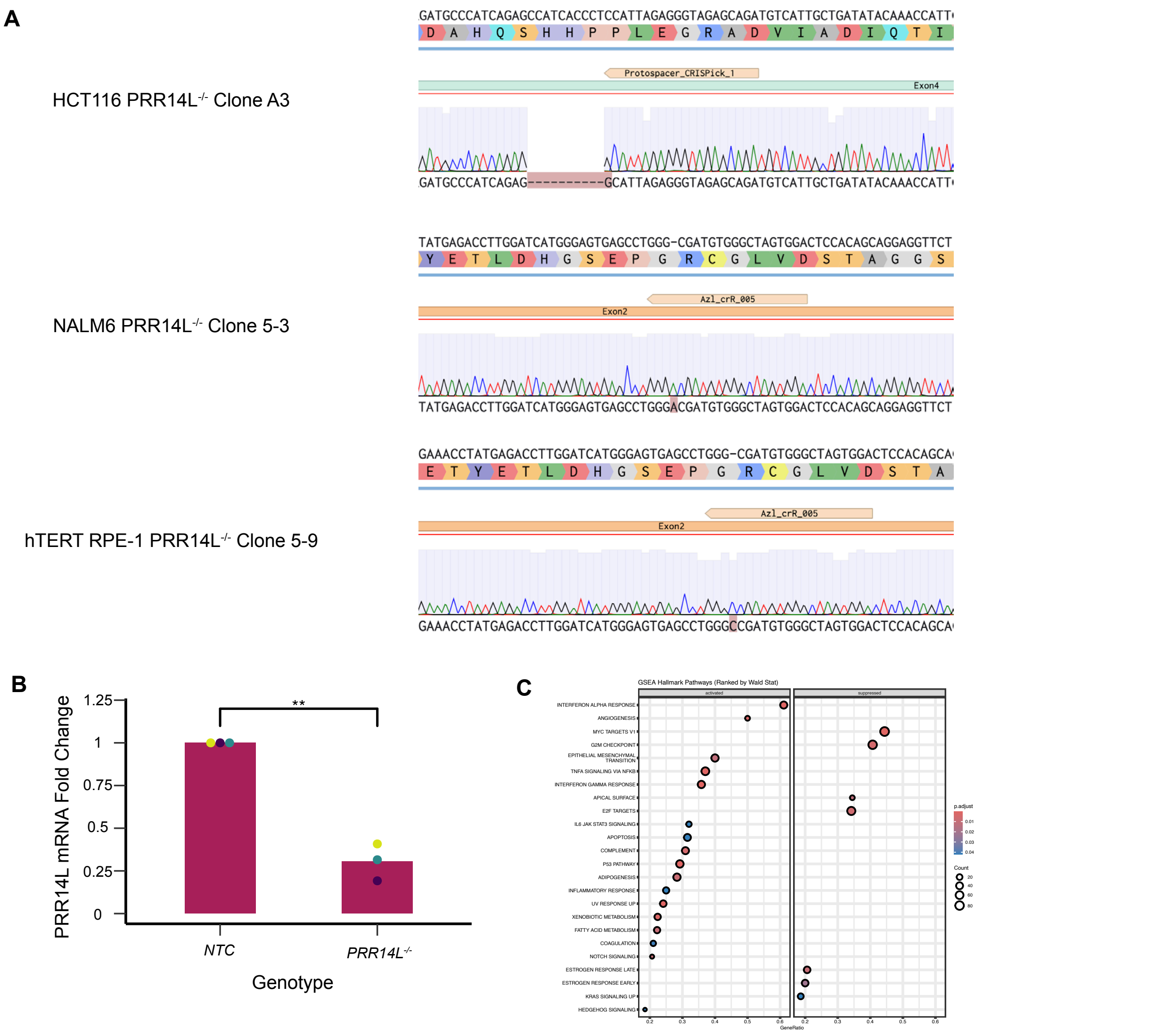


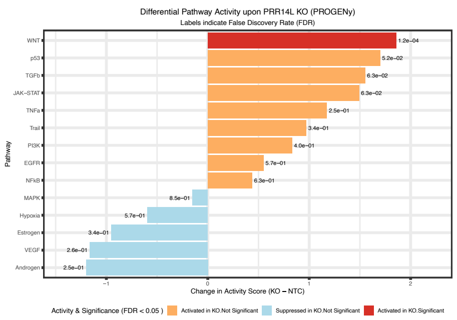


#### Supplemental Figure 2

#### Supplemental Figure 3

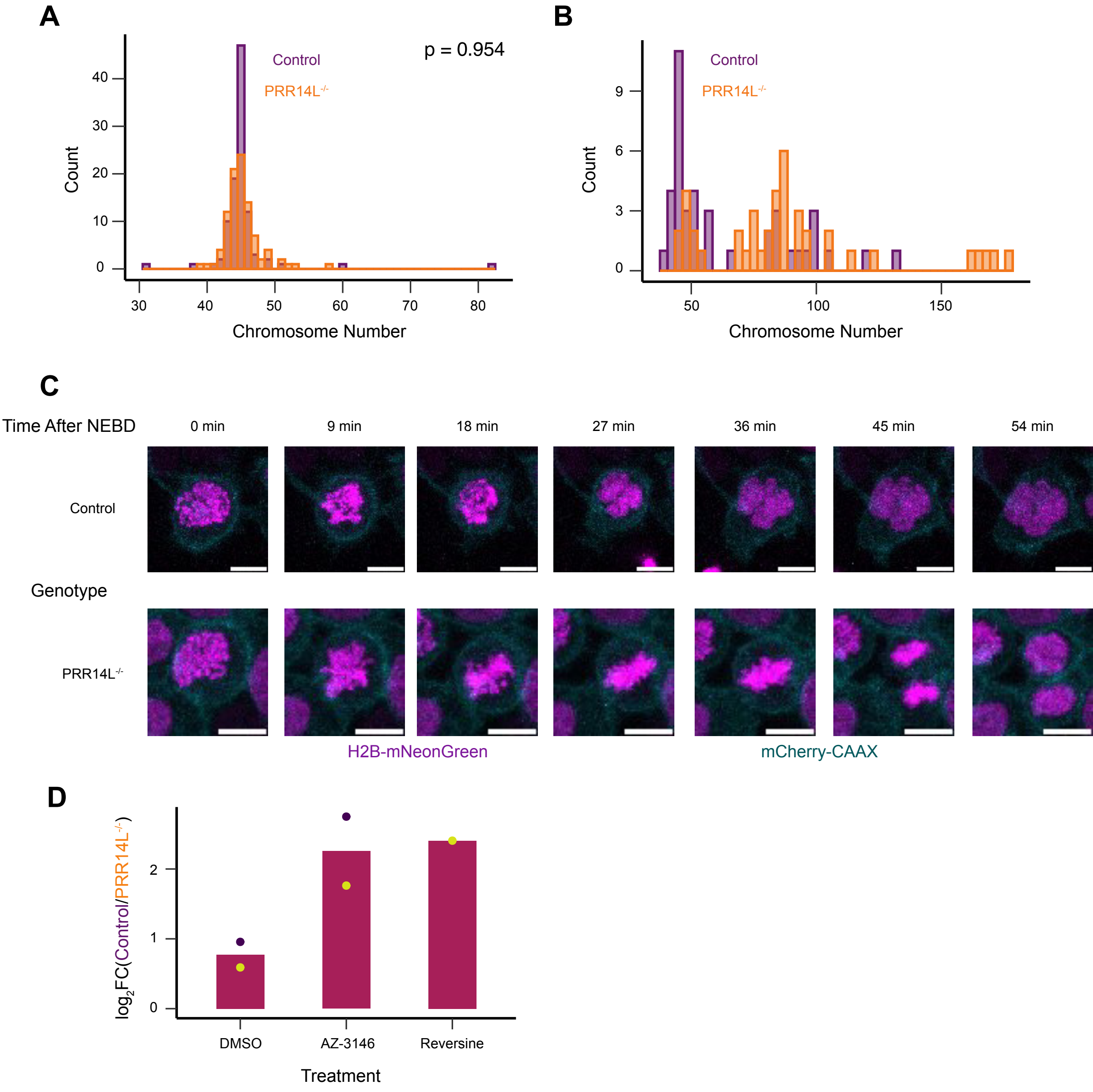
